# Amino Acid-Dependent Material Properties of Tetrapeptide Condensates

**DOI:** 10.1101/2024.05.14.594233

**Authors:** Yi Zhang, Ramesh Prasad, Siyuan Su, Daesung Lee, Huan-Xiang Zhou

## Abstract

Condensates formed by intrinsically disordered proteins mediate a myriad of cellular processes and are linked to pathological conditions including neurodegeneration. Rules of how different types of amino acids (e.g., π-π pairs) dictate the physical properties of biomolecular condensates are emerging, but our understanding of the roles of different amino acids is far from complete. Here we studied condensates formed by tetrapeptides of the form XXssXX, where X is an amino acid and ss represents a disulfide bond along the backbone. Eight peptides form four types of condensates at different concentrations and pH values: droplets (X = F, L, M, P, V, A); amorphous dense liquids (X = L, M, P, V, A); amorphous aggregates (X = W), and gels (X = I, V, A). The peptides exhibit enormous differences in phase equilibrium and material properties, including a 368-fold range in the threshold concentration for phase separation and a 3856-fold range in viscosity. All-atom molecular dynamics simulations provide physical explanations of these results. The present work also reveals widespread critical behaviors, including critical slowing down manifested by the formation of amorphous dense liquids and critical scaling obeyed by fusion speed, with broad implications for condensate function.

## Introduction

Biomolecular condensates are formed via phase separation that is driven by various types of intermolecular attraction, including charge-charge, cation-π, π-π, hydrophobic, and hydrogen-bonding interactions ^1^. Many studies of the phase separation of oppositely charged intrinsically disordered proteins (IDPs) and of basic IDPs with nucleic acids and ATP have highlighted the roles of charge-charge and cation-π interactions ^2-11^. Likewise, cation-π and π-π interactions are found to drive the phase separation of many other proteins ^12-16^. While 1,6-hexanediol has been widely used to disrupt hydrophobic interactions and isolate their contributions to phase separation, urea has been used analogously to probe hydrogen bonds ^17, 18^. Coarse-grained simulations are beginning to predict well the phase-separation threshold concentrations and critical temperatures of IDPs ^19, 20^.

Biomolecular condensates can be found in different material states. A simple way to identify material states is to observe condensate morphologies under a microscope. Spherical droplets are usually assumed to be liquid-like, whereas amorphous aggregates and gels are assumed to be more solid-like. What determines condensate morphologies is still an open question. One can also characterize the material states by measuring material properties, such as the fusion speed of droplets ^13, 21-23^. Slowed or stalled fusion is a sign that condensates are becoming solid-like. Relative to phase-equilibrium properties, our understanding of the determinants of condensate material properties is lagging ^1, 7, 24-26^.

Short peptides have been used to isolate the roles of different amino acids in determining phase equilibrium and condensate morphology. Yuan et al. ^27^ observed the transition of aromatic group-blocked amino acids and dipeptide (b-X, X = A and H; b-XX, X= F) from liquid droplets to nanofibrils over a few hours. Baruch Leshem et al. ^28^ studied W, F, and R-containing peptides (10-14 residues) and identified π-π, cation-π, and hydrogen bonding interactions by Raman and NMR spectroscopy. NMR analysis produced diffusion constants ∼20 Å^2^/ns in the dilute phase; fluorescence recovery after photobleaching (FRAP) indicated a 10,000-fold slowdown in diffusion in the dense phase. Poudyal et al. ^18^ studied the phase separation of homopeptides (X_10_, X = G and V) under crowding by PEG-8000. 20% (w/v) 1,6-hexanediol significantly suppressed the phase separation of V_10_ but 2 M urea had little effect, indicating hydrophobic interactions as the driver for phase separation. The opposite was observed for the effects of additives on G_10_, suggesting that hydrogen bonding plays a major role in its phase separation. Abbas et al. ^17^ synthesized peptides of the form XYssYX, where X, Y = W, F, and L, and ss represents the linker NH-CH_2_-CH_2_-S-S-CH_2_-CH_2_-NH. W-containing peptides formed aggregates whereas F, L-only peptides formed droplets. FFssFF droplets exhibited an inverse fusion speed of ∼1 s/μm and showed robust FRAP over 250 sec. In a follow-up study, these authors showed that YFssFY formed gels containing microstructures that resemble sea urchins ^29^. Over a few hours, FFssFF transitioned from droplets to fibrils that are stabilized by π-π and hydrogen-bonding interactions ^30^.

Here we report the physical properties of condensates formed by peptides of form XXssXX, where X is extended to all the nonpolar amino acids (Figs. S1 and 1a inset). Using brightfield, confocal, and negative-staining electron microscopy, we show that the condensates exhibit a variety of morphologies, including a form, referred to as amorphous dense liquid, that does not appear to have been reported previously. Using optical tweezers, we measured the material properties of these condensates, which differ widely among the peptides, including a 736-fold range in fusion speed and a 3856-fold range in viscosity. Our all-atom molecular dynamics simulations provide physical explanations of these results. Importantly, the peptide condensates display widespread critical behaviors, including critical slowing down manifested by the formation of amorphous dense liquids and critical scaling obeyed by fusion speed.

## Results

### Tetrapeptides form a variety of condensates, including an amorphous dense liquid phase

All eight tetrapeptides (Figs. S1) form condensates, but the condensates have different morphologies when observed under a brightfield microscope (Fig. 1a-f). At high pH (pH 13), condensates with X = F, L, and M are all spherical liquid droplets. The AAssAA condensate is a distinct phase that we refer to as amorphous dense liquid (ADL): amorphous for irregular shapes and dense liquid for the image contrast from the surrounding bulk phase. A small number of droplets may also be mixed in (one indicated by a red arrow). The WWssWW condensate is an amorphous aggregate phase, while the IIssII condensate is a gel phase. Liquid droplets, amorphous aggregates, and gels have been widely reported for biomolecular condensates. ADLs have the appearance of clouds, with a similar morphology to amorphous aggregates but a much lower image contrast. Gels exhibit filamentous features resembling sea urchins, and contrary to the limited size of amorphous aggregates, gel networks can grow indefinitely to span the entire field of view (281.6 μm × 211.2 μm). ADLs were observed as precursors in the formation of crystals ^31^, but do not seem to have been reported previously for biomolecular condensates. Further characterizations of ADLs are presented below.

**Fig. 1.**
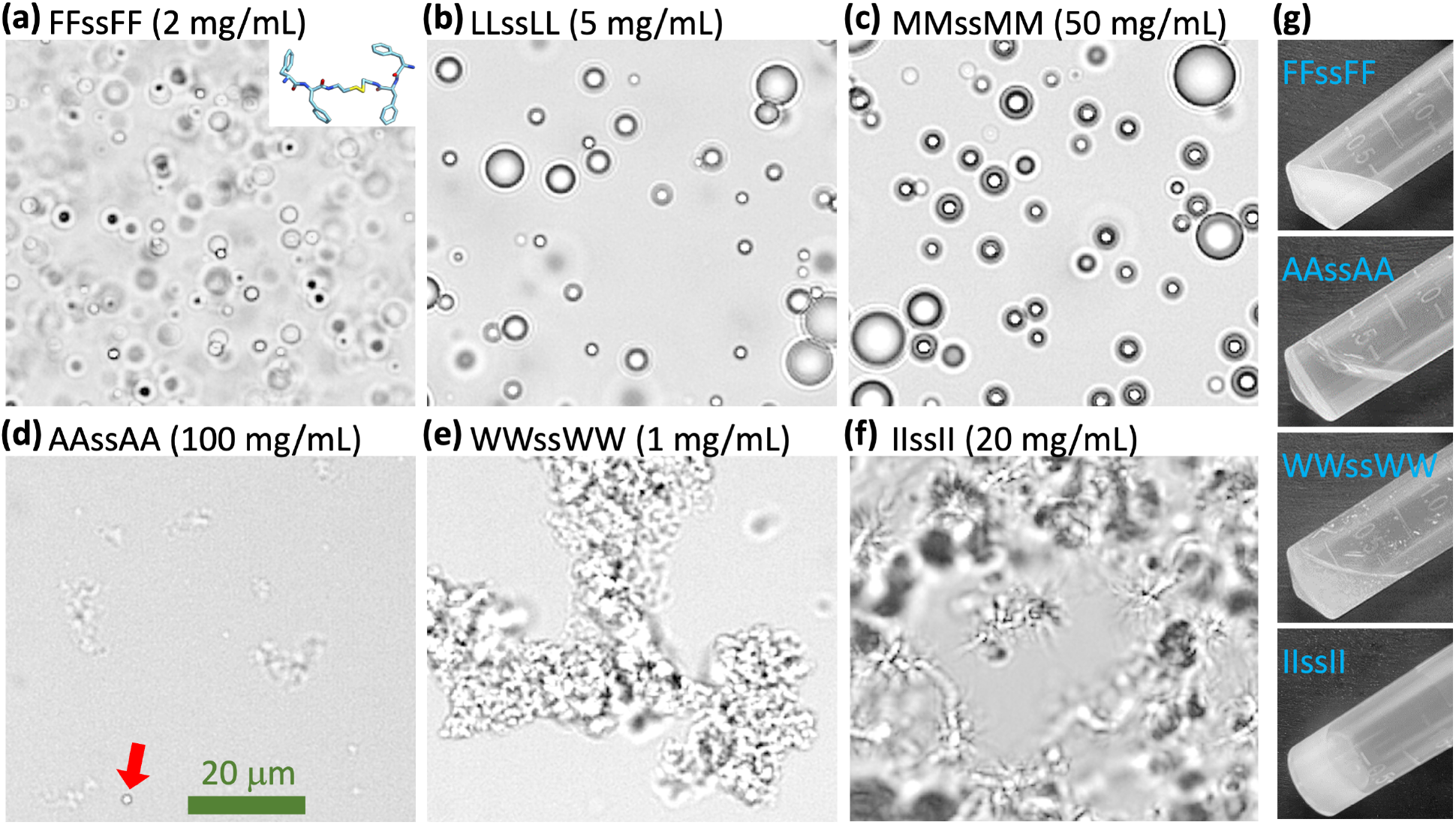
Images showing various forms of tetrapeptide condensates. (a-f) Brightfield images of six tetrapeptide condensates formed at the indicated concentrations in milli Q water and pH 13. In (d), a droplet is indicated by a red arrow. (g) Images of 300-μL samples in a tube, all taken within 2 min of raising pH from 2 to 13.

One way to further distinguish the four different forms of tetrapeptide condensates is to observe them in a tilted test tube with the naked eye (Fig. 1g). Droplet, ADL, and aggregate samples are all liquid-like in the sense that the surface of the solution remains horizontal when the test tube is tilted. In each of these three forms, the individual condensate particles are invisible to the naked eye; the surrounding bulk phase explains the horizontal surface of the solution in a tilted test tube. Droplet samples, as illustrated by FFssFF prepared at 2 mg/mL, have a milky appearance. ADL samples, as represented by AAssAA at 100 mg/mL, are transparent and look almost like a homogenous solution, consistent with the low contrast in the brightfield image (Fig. 1d). For amorphous aggregates, as shown by WWssWW at 1 mg/mL, sedimentation ensues quickly so that the top of the solution becomes transparent while the bottom remains turbid; the tendency of sedimentation can be attributed to denser packing (relative to droplets). In contrast to these liquid-like samples, gel samples are solid-like. At lower concentrations, gels are visible as suspended, turbid particles in the test tube; at higher concentrations, the entire sample is turbid, suggesting that the gels become a single densely connected mesh that sticks to the test tube wall. The sample does not move relative to the tilted test tube, making its surface slanted (Fig. 1g).

The brightfield images in Fig. 1 show the contrast order ADL < droplet < aggregate. This order is confirmed by negative-stain electron microscopy (EM) (Fig. 2a). At pH 13, VVssVV starts to form gels at 5 mg/mL, as indicated by the filamentous appearance in a brightfield image. EM images further demonstrated the filamentous feature. The condensate morphologies of the eight peptides are summarized in Fig. 2b.

**Fig. 2.**
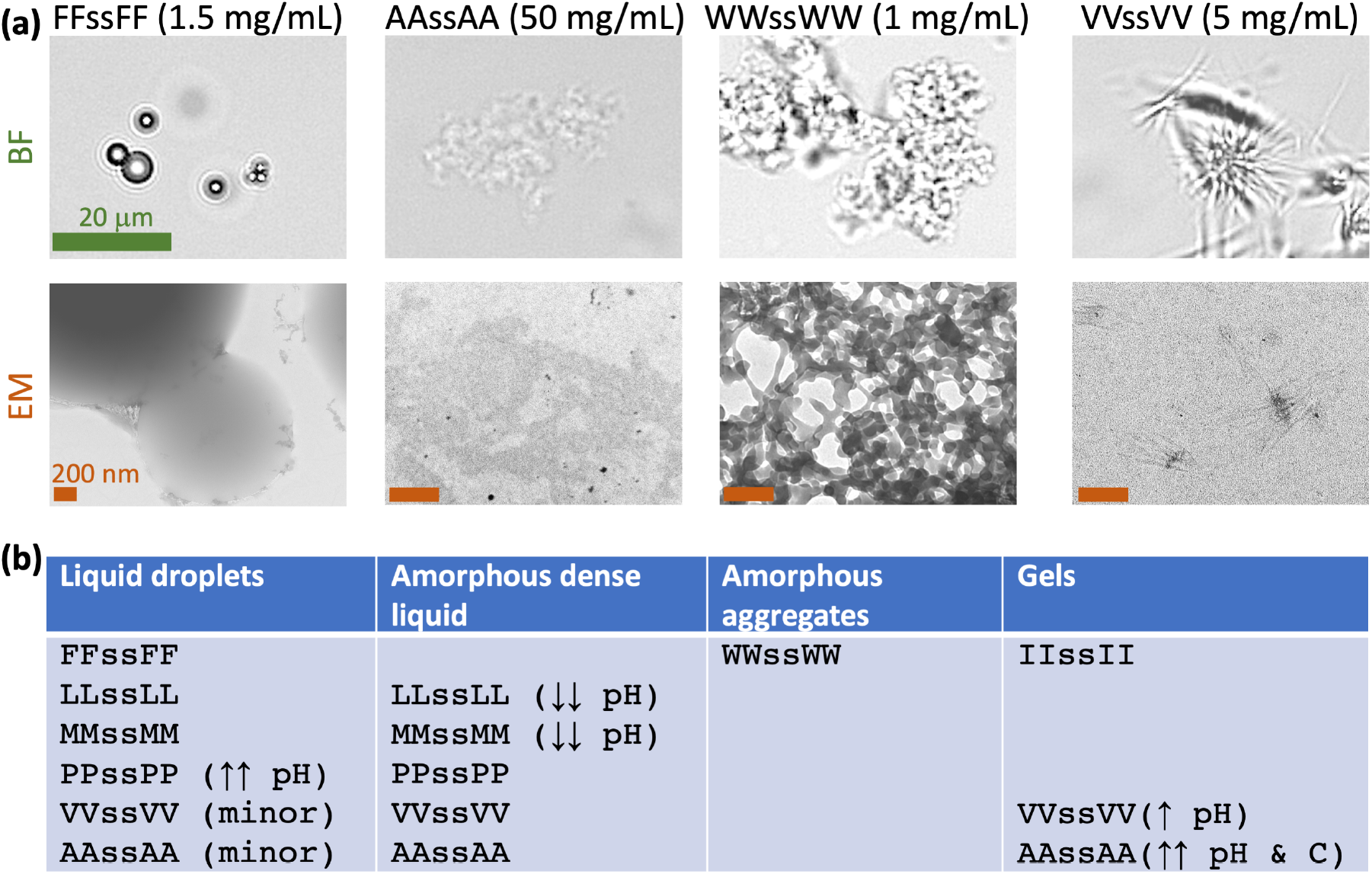
Four forms of condensates formed by tetrapeptides. (a) Contrasting the four forms of condensates by brightfield (BF) and negative-stain electron microscopy (EM). Samples were prepared at the indicated concentrations in milli Q water and pH 13. The green scale bar applies to all the BF images; all the brown scale bars represent 200 nm. (b) Summary of tetrapeptides that form each form of condensate. Single arrows mean roughly one-half of the phase-separation pH range; double arrows mean a small fraction of the phase-separation pH or concentration range.

As one more way to distinguish the four condensate morphologies, we stained the samples with a viscosity-sensitive dye (Fig. S2); the fluorescence intensity provides a measure of the local peptide density. As expected, the fluorescence intensity is very uniform inside FFssFF droplets. It is also uniform inside WWssWW aggregates, such that regions showing intense fluorescence match those that are in focus in the corresponding brightfield image; these regions have rough boundaries instead of the smooth, circular ones of droplets. By contrast, the dye appears to stain only dense cores in AAssAA ADLs and IIssII gels. These core regions apparently are tiny for ADLs but more spread out for gels.

We used FRAP to probe molecular mobility inside condensates (Fig. S3). A 200 nm × 200 nm square region inside condensates was photobleached; the subsequent recovery was fit to an exponential function to obtain the mobile fraction of dye molecules. This fraction is 61% inside FFssFF droplets (comparable to what was reported by Abbas et al. ^17^) and reduces to 29% inside WWssWW aggregates and only 18% inside IIssII gels. The latter result suggests that the gel networks are largely solid-like. Like droplets and ADLs, WWssWW aggregates form instantaneously but IIssII gels take minutes to fully form when started inside the phase-separation region but close to the phase boundary.

The morphology results summarized in Fig. 2b were based on the phase diagrams of the peptides, determined by scanning pH and peptide concentration (*C*) and observing the sample at the given pH and *C* under a brightfield microscope. We dissolved the peptides in 50 mM imidazole buffer to better tune the pH, at least for the pH 6.2 – 7.8 operating range of this buffer. The outcome is either no phase separation (“×” in Fig. 3a-e) or, when phase separation occurs, the condensate morphology (“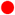” for droplets, “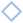” for ADLs, and “✲” for gels). Only a single condensate morphology is observed for three peptides: WWssWW as amorphous aggregates, IIssII as gels, and FFssFF as droplets (Fig. 3a). For LLssLL (Fig. 3b) and MMssMM (Fig. 3c), droplets are observed in most of the phase-separated region of the phase diagram, except at low pH, where ADLs are formed; the opposite is true of PPssPP. For VVssVV (Fig. 3d), the phase-separated region is divided into a gel sub-region (higher pH) and an ADL sub-region (lower pH). AAssAA is similar (Fig. 3e), except that its gel sub-region is limited to very high pH and very high peptide concentration.

**Fig. 3.**
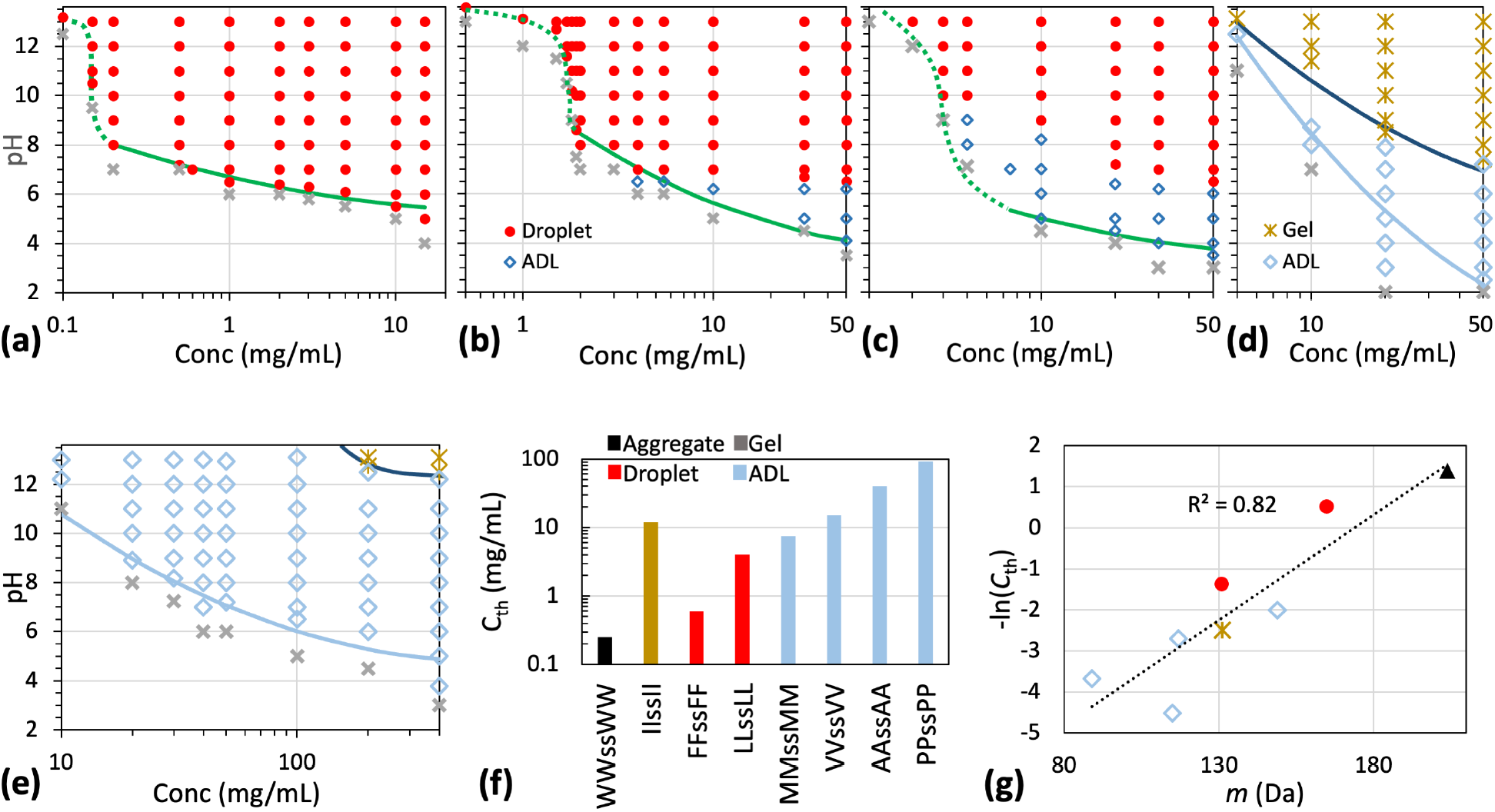
Phase diagrams and threshold concentrations at pH 7. (a-e) Phase diagrams of tetrapeptides with X = F, L, M, V, and A in 50 mM imidazole buffer. For LLssLL and MMssMM, the transition from ADLs to droplets spans 1 unit of pH values; in the transition region, the morphology is assigned to the dominant species. (f) Threshold concentrations for phase separation at pH 7. (g) Correlation between threshold concentration and amino-acid molecular mass.

### Phase-separation threshold concentration correlates with sidechain interaction strength

To compare the drive for phase separation among the peptides, we measured the minimum, or threshold, concentrations (*C*_th_) for phase separation at pH 7 (Fig. 3f). The values of *C*_th_ range from 0.25 mg/mL for WWssWW to 92 mg/mL for PPssPP. This 368-fold difference demonstrates the enormous disparity between amino acids in their drive for phase separation. Because the amino acids in all the eight homopeptides are nonpolar and the strengths of sidechain-sidechain interactions are expected to scale with the amino-acid sizes, we tested for correlation between –ln(*C*_th_) and the amino-acid molecular mass *m* (Fig. 3g). There is a strong correlation between these two properties, with the coefficient of determination (*R*^2^) at 0.82. A slightly higher correlation, *R*^2^ = 0.89, is obtained with a compound factor, *σ*^3^*λ*, where *σ* is the Lennard-Jones diameter and *λ* is the “stickiness” parameter of a coarse-grained model for IDPs (one-bead per amino acid) ^19^. This compound factor was devised to capture the strengths of nonpolar interactions between amino acids ^1^.

We wondered whether backbone hydrogen bonding also contributed to the drive for phase separation. Urea was used as a reporter for backbone hydrogen bonding in previous studies ^17, 18^. Consistent with the observation of Abbas et al. ^17^, urea has a modest effect on droplet formation of 0.25 mg/mL FFssFF at pH 13 (Fig. S4, first column). In contrast, droplet formation of 1.5 mg/mL LLssLL is inhibited by 2 M urea (Fig. S4, second column). Likewise, gel formation of 1.5 mg/mL IIssII is greatly suppressed by 2 M urea and blocked by 3 M urea (Fig. S4, third column). These observations suggest that backbone hydrogen bonding plays an appreciable role in the phase separation of LLssLL and IIssII.

The boundary of the phase-separated region moves to higher peptide concentrations as the pH is lowered. pH determines the charge states of the terminal amines of each peptide and, hence, the extent of net charge repulsion between peptide molecules. Peptide concentration controls the density of intermolecular interactions. Apparently, greater net charge repulsion must be balanced by a higher density of interactions between amino acids for phase separation to occur. The phase boundary can be fit to a parabolic function:

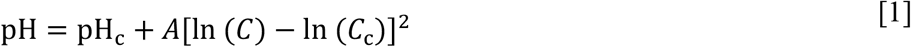

where pH_c_ is the minimum or critical pH at which phase separation is still observed, *C*_c_ is the corresponding critical concentration, and *A* is a constant. The critical pH values are 5.3, 3.8, 3.5, 0.3, and 4.8 for peptides with X = F, L, M, V, and A, respectively. The corresponding critical concentrations are 66, 148, 154, 245, and 665 mg/mL, respectively. These critical values can only be taken as qualitative indications as we are limited by the highest concentrations for phase-boundary determination. The phase boundaries of peptides with X = F, L, and M appear to show a transition above pH 8, with the caveat that the pH regulation capacity of the imidazole buffer is weak in this range.

At pH 7 and the respective threshold concentrations, the condensates formed are amorphous aggregates (X = W), gels (X = I), liquid droplets (X = F and L), or ADLs (X = M, P, V, and A) (Figs. S5 and 3f). While the threshold concentrations correlate well with the strengths of sidechain nonpolar interactions, the rules governing condensate morphology are more complex. Sidechain interaction strength appears to explain three of the four morphologies (Fig. 3f), as amorphous aggregates, droplets, and ADLs are favored by stronger (X = W), intermediate (X = F and L), and weaker (X = M, P, V, and A) interactions, respectively. This is also the contrast order exhibited by brightfield and EM images of the condensates. However, it is difficult to predict when gels will form. For example, L and I are very similar chemically, but LLssLL favors droplets whereas IIssII only forms gels.

### ADLs may be a manifestation of critical slowing down

In all the cases where ADLs are observed, they occur near the critical point (Fig. 3b-e and S5). ADLs often are mixed with a small number of droplets (Figs. 1d and S5). As we move away from the critical point, more and more droplets are formed, to the point where only droplets are found in the case of X = L, M, and P (Fig. 3b-c and S5). Together, these facts suggest that ADLs are metastable precursors to droplets, and their occurrence is due to critical slowing down. The usual explanation for critical slowing down is that, near the critical point, the correlation length increases indefinitely, and correspondingly, the characteristic relaxation time for phase transition grows to infinity ^32^. It looks like this explanation applies to the slow conversion of ADLs into droplets.

To directly explore the conversion from ADLs to droplets, we asked whether raising the temperature could speed up the conversion. Abbas et al. ^17^ already showed that the phase separation of XXssXX peptides had an upper critical solution temperature (UCST), meaning that raising the temperature decreases the drive for phase separation. We confirmed the UCST character by, e.g., a decreasing number of FFssFF droplets as the temperature was raised from 4 °Cto 25 and 100 °C(Fig. S6, column 1). Still, we reasoned that a high temperature, by enhancing thermal fluctuations, could speed up the kinetics of phase transition. We checked the temperature dependence of the condensate morphologies of peptides with X = L (50 mg/mL and pH 6.2), V (50 mg/mL and pH 7.2), and A (100 mg/mL and pH 13) (Fig. S6, columns 2-4).

Whereas droplets are rare to find among ADLs at 4 and 25 °C, they are readily seen along with ADLs at 100 °C. Once formed as droplets at 100 °C, they remain as droplets even when the samples are cooled down to 25 °C, supporting the notion that ADLs are metastable precursors to droplets.

### Droplet fusion speed exhibits critical scaling with pH

Next we report the material properties of tetrapeptide droplets, including fusion speed, interfacial tension (*γ*), and zero-shear viscosity (*η*), all measured on a LUMICKS dual-trap optical tweezers (OT) instrument. OT-directed fusion was set up by trapping two equal-sized droplets and bringing them into contact (Fig. 4a) ^23, 33^. The subsequent spontaneous fusion process was monitored by the small force signal due to the retraction of the droplets from the fixed traps. The force trace was fit to a stretched exponential to obtain the fusion time *τ*_fu_ (Fig. 4b). For LLssLL droplets of similar sizes (initial radius *R* ∼ 3 μm), the fusion time has a clear pH dependence, increasing with decreasing pH (Fig. 4b). By fitting the dependence of *τ*_fu_ on *R* to a proportional relation, we obtained inverse fusion speed as the slope, *τ*_fu_/*R* (Figs. 4c and S7). We verified that the initial peptide concentration did not affect the fusion speed (Fig. S7b).

**Fig. 4.**
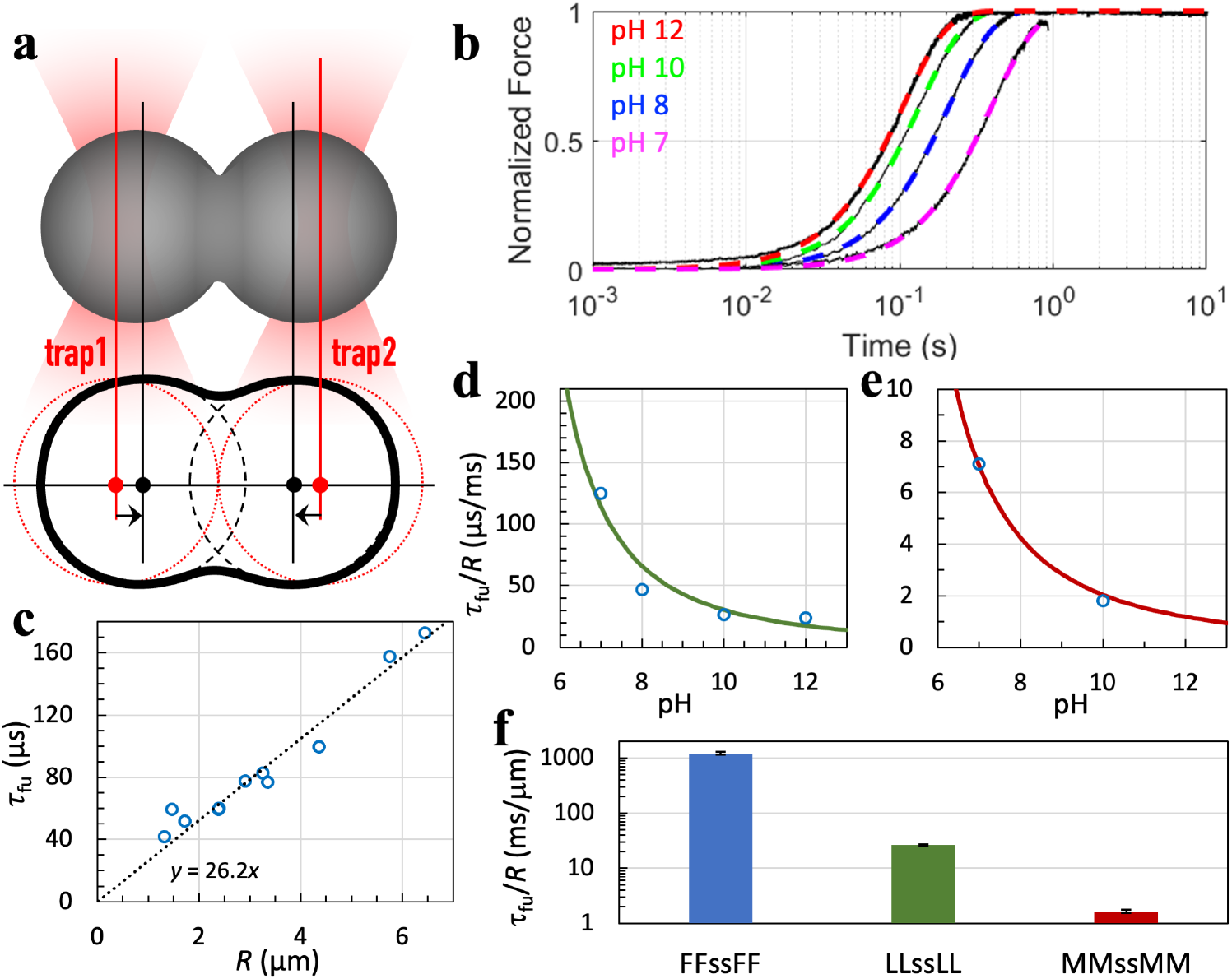
OT-directed droplet fusion. (a) Illustration of directed fusion. (b) Fusion progress curves of LLssLL droplets. Black traces show raw data; colored curves are fits to Eq [9]. Droplets were prepared at the indicated pH and grown to 3 to 4 μm in radius before fusion. (c) Fusion time of LLssLL droplets as a function of initial droplet radius. Each circle represents a fusion event. A line displays the proportional relation between fusion time and droplet radius. Droplets were prepared with 5.5 mg/mL LLssLL at pH 10. (d) Inverse fusion speed of LLssLL vs. pH. The curve displays a fit to Eq [2] with *α* fixed 2, pH_c_ fixed at 3.8, and *b* adjusted to 1165. (e) Inverse fusion speed of MMssMM vs. pH. The curve displays a fit to Eq [2] with *α* fixed 2, pH_c_ fixed at 3.5, and *b* adjusted to 86. (f) Inverse fusion speeds of droplets with X = F, L, and M at pH 10. Error bars represent the standard error in *τ*_fu_/*R* as a fitting parameter.

The effect of pH on *τ*_fu_/*R* can be modeled by a scaling law,

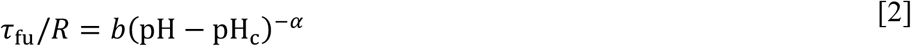

where *α* is a critical exponent and *b* is a constant. For purely viscous droplets, one expects ^23^

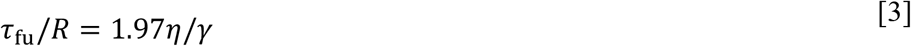

*γ* obeys the scaling law

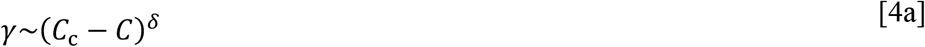

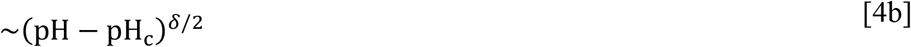

where *δ* ≈ 4 ^34^ and Eq [1] has been used on going from the first line to the second line. *η* does not exhibit critical singularity (i.e., it remains finite at the critical point). Eqs [3] to [4b] predict *α* = *δ*/2 ≈ 2. For both LLssLL and MMssMM droplets, the pH dependence of *τ*_fu_/*R* follows the scaling law with the critical exponent *α* = 2 (Figs. 4d, e). However, although Eq [3] serves as a motivation for the scaling law for *τ*_fu_/*R*, it breaks down for the peptide condensates studied here, as shown below, because of their viscoelastic nature ^11, 24^.

### Fusion speeds of tetrapeptide droplets span three orders of magnitude

We compare the inverse fusion speeds of droplets with X = F, L, and M in Fig. 4f. For this comparison as well as comparisons of other material properties below, we selected pH 10 because there the peptides readily form a large number of droplets. MMssMM droplets fuse at *τ*_fu_/*R* = 1.7 ± 0.1 ms/μm. The corresponding fusion speed is 16-fold faster than that of LLssLL droplets and 736-fold faster than that of FFssFF droplets. (The inverse fusion speed measured here is comparable to that reported by Abbas et al. ^17^.) The fusion speeds follow the same order as the threshold concentrations (Fig. 3a-c), suggesting sidechain interaction strength as a determinant for fusion speed. However, whereas the threshold concentrations differ by only 20-fold, the fusion speeds differ by 736-fold.

At pH 10, PPssPP forms a significant portion of ADL along with droplets, making it difficult to study droplet fusion. We thus measured the fusion speed at pH 13 (Fig. S7f), where only droplets are formed. The inverse fusion speed is 0.20 ± 0.01 ms/μm. When the scaling law is used to predict an inverse fusion speed for MMssMM, a value of 0.95 ms/μm is obtained. The latter value indicates that PPssPP has a higher fusion speed than MMssMM, which is consistent with the order in threshold concentration between these two peptides (Fig. 3f) and further supports sidechain interaction strength as a determinant for fusion speed.

### Interfacial tensions fall into a relatively narrow range

As Eq [3] indicates, droplet fusion is driven by interfacial tension and retarded by viscosity. By using two optically trapped beads to stretch a droplet (Fig. 5a) and monitoring the forces at successive extensions (Fig. 5b), we measured the interfacial tension ^24, 35^. For droplets formed by peptides with X = F, L, and M, the interfacial tensions are 96 ± 9, 109 ± 16, and 38 ± 3 pN/μm (Fig. 5c). The values differ by only 3-fold. Interfacial tensions of various biomolecular condensates all fall into a relatively narrow range, mostly from 20 to 200 pN/μm ^1^.

**Fig. 5.**
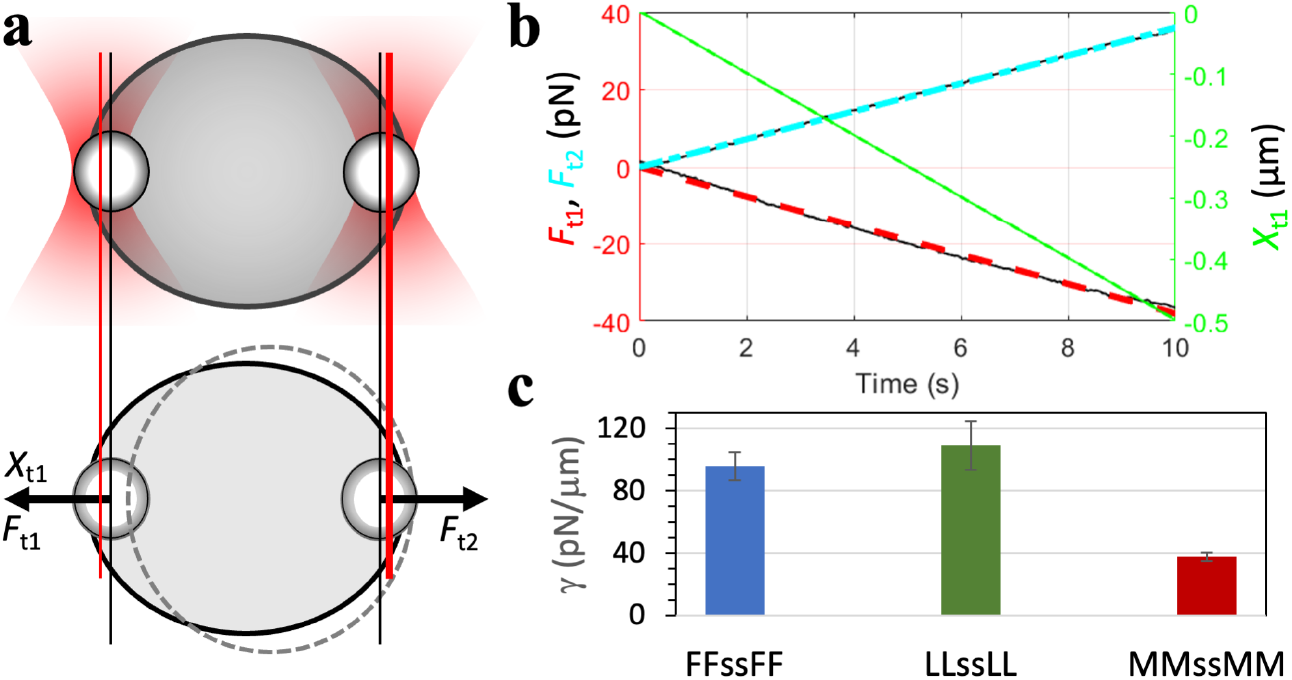
Interfacial tensions of tetrapeptide droplets. (a) Stretching of a droplet by two optically trapped beads (1.13 μm in radius). (b) Stretching forces and extensions measured on an LLssLL droplet formed at 5.5 mg/mL and pH 10. (c) Interfacial tensions of tetrapeptide droplets with X = F, L, and M at pH 10. Error bars represent standard deviations among four droplets with ∼6 μm radii.

### Effective viscosity during droplet fusion is much lower than zero-shear viscosity

With the fusion speed and interfacial tension at hand, we can deduce the effective viscosity, *η*_eff_, in the fusion process. *η*_eff_ is defined by inverting Eq [3]:

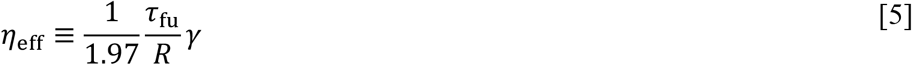

The values for droplets formed by peptides with X = F, L, and M are 59, 1.5, and 0.032 Pa s. IDP condensates have a tendency to exhibit effective viscosities much lower than their zero-shear viscosities, a phenomenon called shear thinning ^1, 24, 36^.

To test whether tetrapeptide droplets also exhibit shear thinning, we measured their zero-shear viscosities using a brightfield camera to track a free bead inside a droplet that was settled on a coverslip (Fig. 6a, b) ^36^. By fitting the mean-square displacement (MSD) of the tracked bead to a linear function of time (Figs. 6c and S8), one can find the zero-shear viscosity *η* from the slope. The results are 856 ± 351, 123 ± 30, and 0.22 ± 0.06 Pa s, respectively, for peptides with X = F, L, and M. These values are 7-to 85-fold higher than the corresponding effective viscosities, thereby implicating substantial shear thinning. The extent of shear thinning exhibited by the tetrapeptides here is much greater than that by IDPs ^24^ but somewhat less than that by ATP-IDP mixtures ^11^. The enormous shear thinning in the latter systems was partly attributed to the small size of ATP, allowing it to quickly reform bridges between IDP chains during droplet fusion. Given the relatively small size of the tetrapeptides, perhaps a similar mechanism is at play here.

**Fig. 6.**
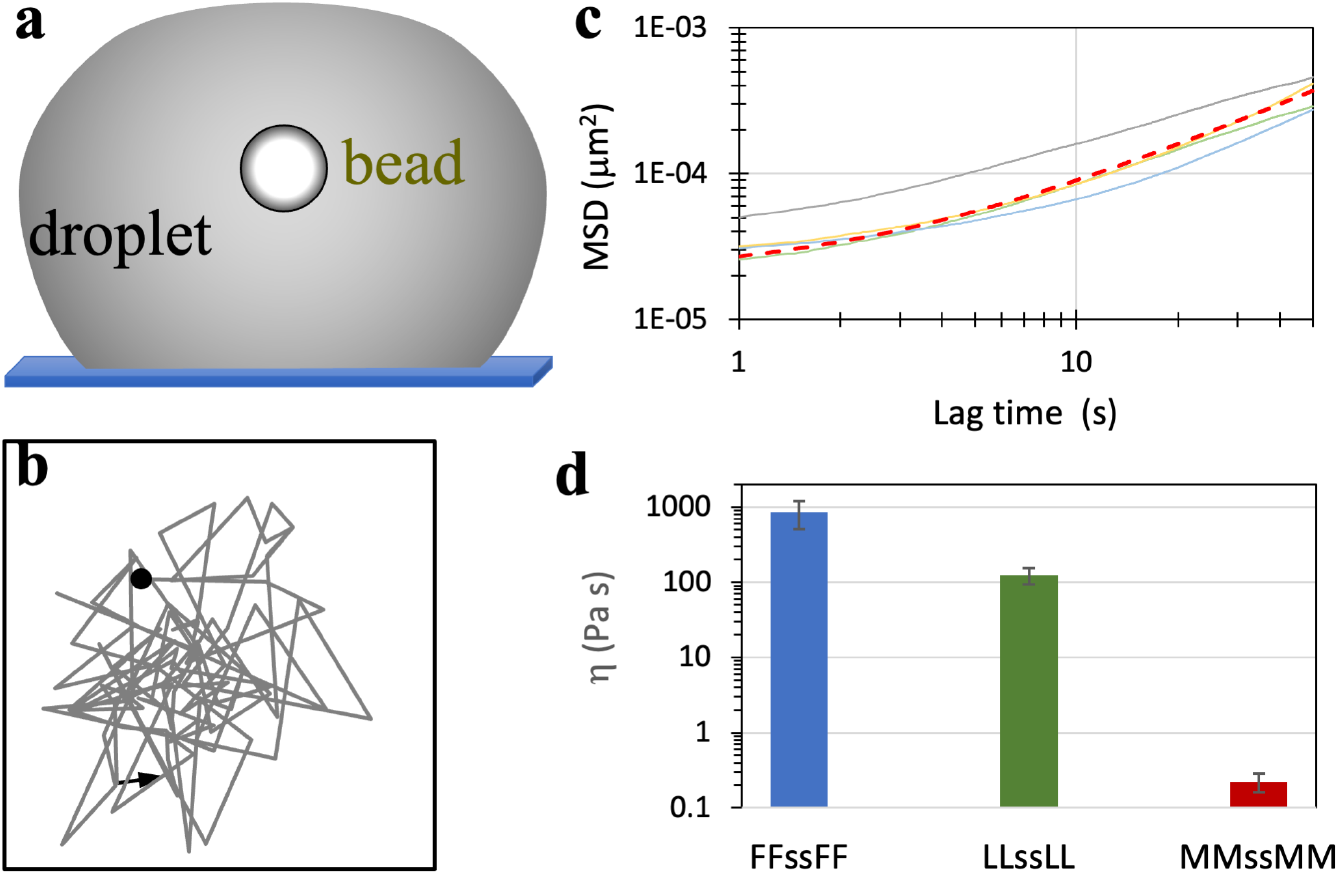
Zero-shear viscosities of tetrapeptide droplets. (a) Tracking of a free bead inside a settled droplet. (b) A short 2-dimensional trajectory of a bead (1.13 μm in radius) in an LLssLL droplet prepared at 5.5 mg/mL and pH 10, tracked by a brightfield camera. (c) MSDs (solid curves) of beads inside four LLssLL droplets. The fit to a linear function for one MSD curve (orange) is displayed as a red dashed curve. (d) Zero-shear viscosities of tetrapeptide droplets with X = F, L, and M at pH 10. Error bars represent standard deviations among four beads.

It is worth noting that the viscosity measured here for the FFssFF condensate appears to be higher than any previously reported value for biomolecular condensates ^1^. It is higher than the LLssLL viscosity by 7-fold and the MMssMM viscosity by nearly 4000-fold.

### Molecular dynamics simulations of tetrapeptides recapitulated condensate morphologies

Previously we observed spontaneous condensate formation in molecular dynamics simulations of 64 copies of FFssFF in explicit water ^37^. Here we changed to a force-field combination that was found to model well IDPs in explicit water ^38^ and expanded the simulations to all eight tetrapeptides. For each tetrapeptide, 64 loosely packed copies of the form with both terminal amines neutral (modeling high pH) were solvated in a cubic simulation box. Four different fractions of water molecules were removed to concentrate the peptide to a range of initial concentrations; duplicate simulations of each of these systems were carried out at constant temperature (294 K) and pressure (1 bar). Within 1 μs of the simulations, condensates start to take shape. For peptides with X = F, M, L, I, and V, the condensates in at least one of the eight simulations are a stable slab (Figs. 7a and S9), which corresponds to a spherical droplet in a bulk solution. The peptide molecules in these condensates exhibit a moderate level of dynamics (Movie S1 for LL), with 20% to 40% of molecules surrounding a tagged molecule moving away from that molecule after 1 μs (Fig. S10a). In comparison, WWssWW condenses only into an amorphous aggregate (Figs. 7a and S9), which remains static except for a few molecules on the surface (Movie S2 for WW; Fig. S10a,b). By contrast, AAssAA and PPssPP also form slabs but the boundaries of the slabs are typically rough (Figs. 7a and S9) and change rapidly. In 1 μs, ∼80% of molecules surrounding a tagged molecule move away from that molecule (Movie S3 for AA; Fig. S10a,c).

**Fig. 7.**
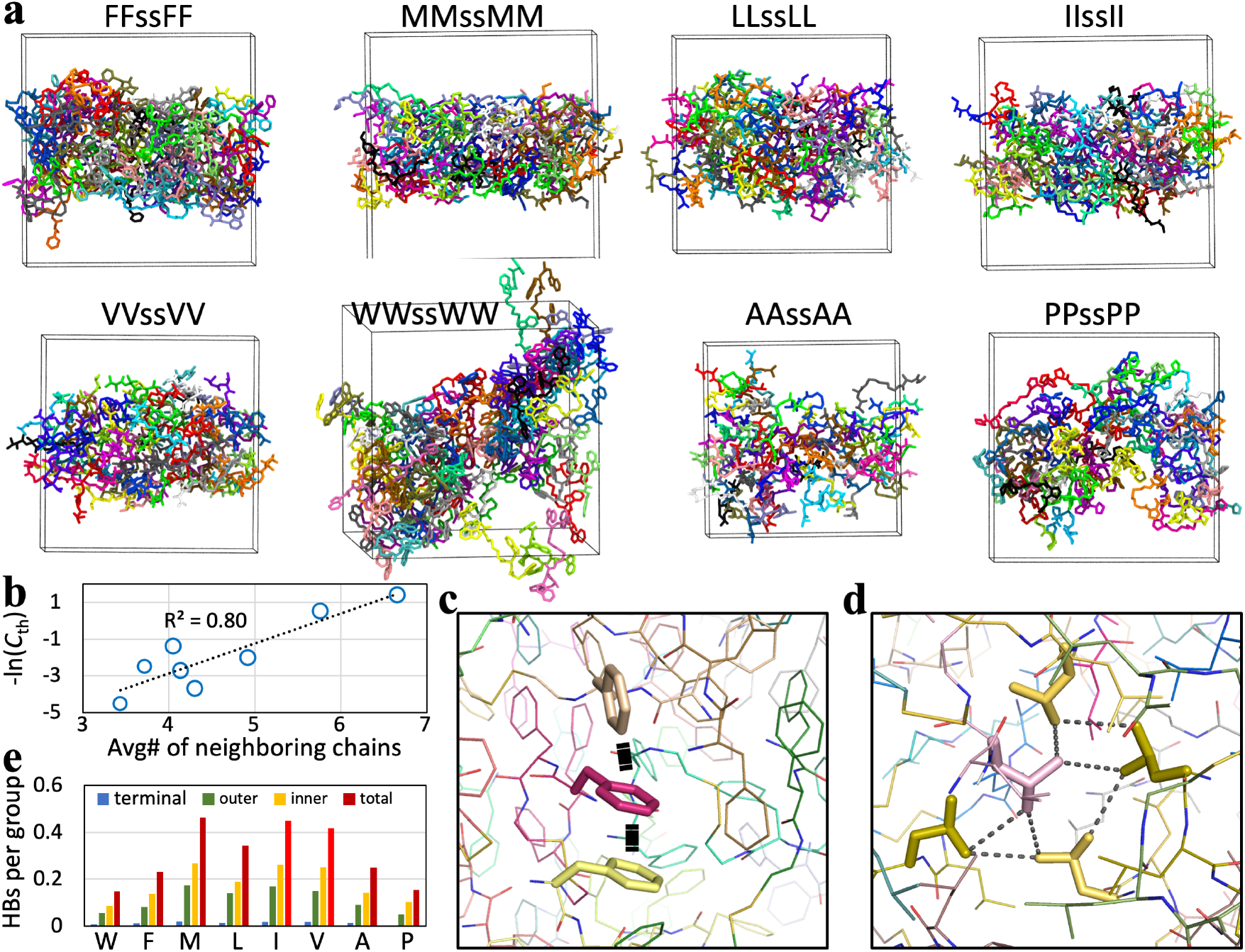
Condensate morphologies and intermolecular interactions in molecular dynamics simulations. (a) Morphologies of the eight tetrapeptides. (b) Correlation of the average number of chain neighbors with phase-separation threshold concentration. (c) π-π interactions in the FFssFF condensate. (d) A hydrophobic cluster in the LLssLL condensate. (e) Average numbers of hydrogen bonds formed by the terminal amine, the outer peptide group, or the inner peptide group.

Whereas peptides with X = F, M, L, A, and P form slabs in four or five of the eight simulations, IIssII and VVssVV do so in only one and two, respectively, of the eight simulations. In the remaining simulations, the condensates have many holes, which perhaps resemble gels (Fig. S11). Overall, the simulations show that peptides with X = F, M, and L favor droplets, peptides with X = A and P form condensates that partly resemble droplets and partly resemble ADLs, peptides with X = I and V prefer gel-like condensates, and WWssWW only forms aggregates. These results match well with the high-pH condensate morphologies observed under a microscope.

The threshold concentration quantifies the propensity to phase separate: a lower *C*_th_ corresponds to a higher propensity. A determinant of this propensity is the ability of the component molecules to network with each other. As a measure of the latter property, we calculated the number of neighboring chains of each peptide molecule in the condensate (Fig. S12). The average number of neighboring chains ranges from 6.7 for X = W to 3.4 for X = P. A strong correlation is found between the average number of neighboring chains and –ln(*C*_th_), with *R*^2^ at 0.80 (Fig. 7b).

Intermolecular networks in the condensates are stabilized by π-π interactions for X = W and F and hydrophobic clusters for the other peptides (Figs. 7c,d and S13). The backbones of the peptides also exhibit varying levels of hydrogen bonding. LLssLL and IIssII condensates have much higher levels of backbone hydrogen bonding than FFssFF condensates, consistent with the experimental results reported by adding urea (Fig. S4). Interestingly, the hydrogen-bonding levels of both IIssII and VVssVV are particularly high, on average forming close to one hydrogen bond per chain. Backbone hydrogen bonding perhaps contributes to gel formation.

To study the equilibration between the dense and dilute phases of FFssFF and LLssLL peptides, we placed the dense slab formed in the simulations with a cubic box to an elongated box and also varied the charge states of the terminal amines to model different pH values (Fig. 8a). The numbers of chains with zero, single, and double charges were determined by assuming p*K*_a_ values of 7 and 6 for the two amines in each chain. For example, at pH 7.3, half of the 64 copies are neutral at both amines and the other half are charged at one amine. With decreasing pH, the dilute-phase concentrations increase significantly (Fig. 8a-c), due to net charge repulsion in the dense phase. The dilute-phase arm of the FFssFF and LLssLL binodals is qualitatively similar to the experimental phase boundaries of Fig. 3a,b. The dilute-phase concentrations from the simulations of FFssFF and LLssLL at pH 7.3 are 2 and 7 mg/mL, respectively, comparable to the experimental *C*_th_ values of 0.5 and 3 mg/mL (Fig. 3f). In comparison, the dense-phase concentrations decrease but only slightly with decreasing pH.

**Fig. 8.**
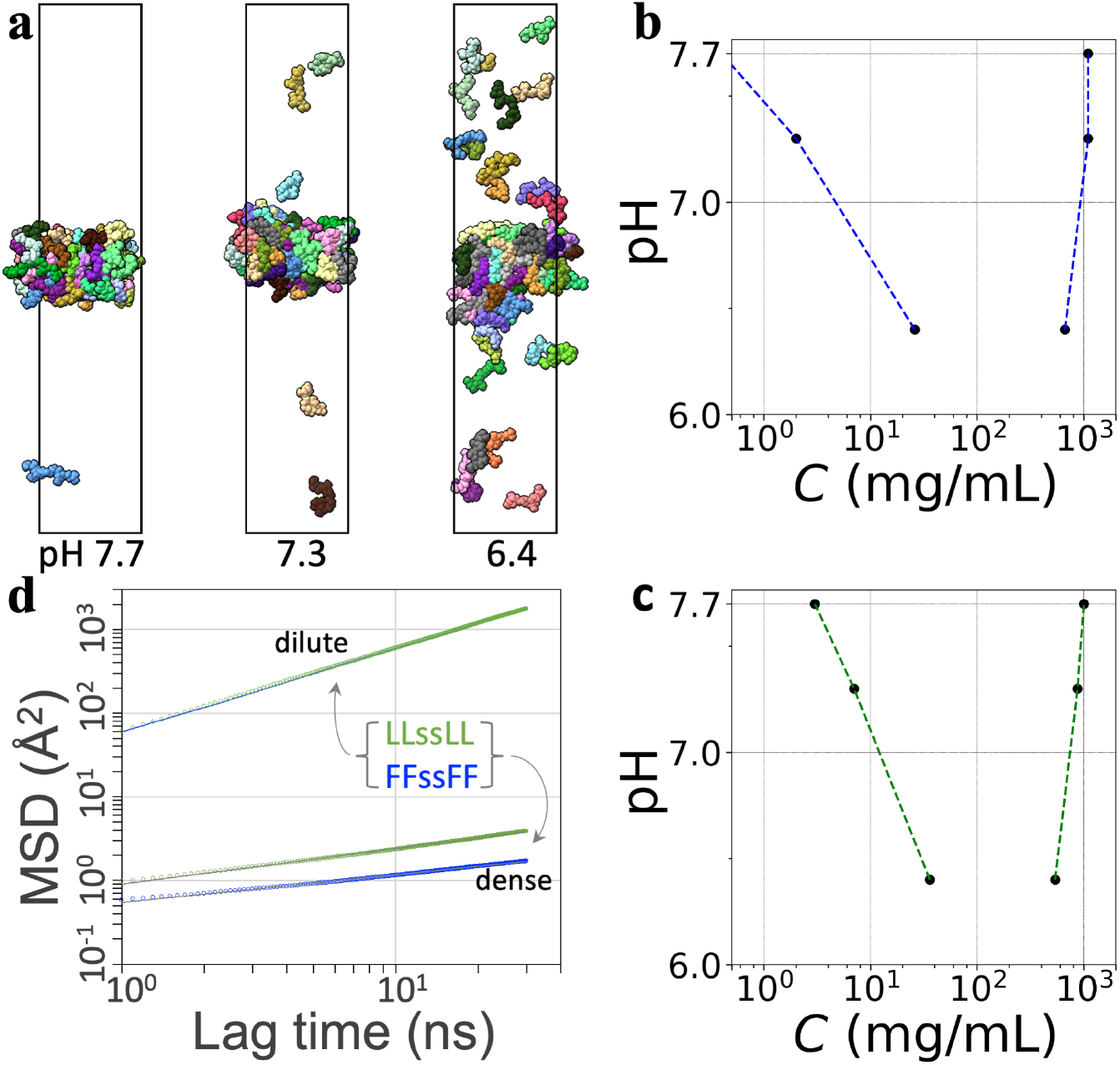
Properties of the dilute and dense phases of FFssFF and LLssLL. (a) Phase coexistence of LLssLL at different pH values. (b) Binodal of FFssFF. (c) Binodal of LLssLL. (d) MSDs of tetrapeptide molecules in the dilute and dense phases at pH 7.3. For the dilute phase, the results for FFssFF and LLssLL are displayed as a blue line and green circles, respectively; for the dense phase, the raw data are displayed as blue or green circles and a fit to Eq [6] is shown as a black line.

To probe dynamics within the condensates, we calculated the diffusion constants of peptide molecules that stayed either in the dilute phase or in the dense phase in a 1.04-μs period (Fig. 8d). In the dilute phase, the MSDs of FFssFF and LLssLL are essentially identical and conformed to a Brownian behavior, with time dependence given by 6*D*_0_*t* and a diffusion constant *D*_0_ = 9.95 Å^2^/ns. This has the same order of magnitude as that for dilute-phase short peptides measured by NMR ^28^. In the dense phase, the MSDs exhibit subdiffusion, which is characteristic of molecular motions hindered by mobile obstacles ^39^. The time dependence of the MSDs fits well to

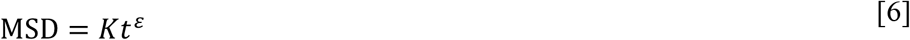

with *ε* = 0.33 for FFssFF and 0.42 for LLssLL. The subdiffusion can be viewed as a manifestation of condensate viscoelasticity at the sub-μs timescale ^40^. We define an effective diffusion coefficient

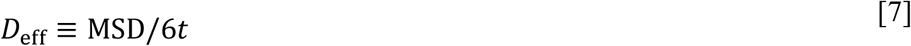

The difference in *D*_eff_ between FFssFF and LLssLL grows with *t* and is 2.3-fold at *t* = 30 ns. The theoretical expectation is that *D*_eff_ would become a constant at much longer times. If Eq [6] is extrapolated to 0.5 ms, it would predict *D*_eff_ = 1.4 × 10^−5^ Å^2^/ns for FFssFF and 7.9 × 10^−5^ Å^2^/ns for LLssLL. The corresponding values of *D*_eff_/*D*_0_ would yield dense-phase viscosities, 700 and 126 Pa s, that match well with the measured values (Fig. 6d), with the caveat that such extrapolation carries significant uncertainty.

Two interrelated factors explain the lower diffusion coefficient and hence higher viscosity of FFssFF relative to LLssLL. First, as noted above, the amino acid F has a greater sidechain interaction strength than L. This difference is reflected by the higher average number of neighboring chains for FFssFF than for LLssLL. Second, as shown in Fig. 8b,c, the FFssFF dense phase has higher densities than the LLssLL counterpart.

The difference in dense-phase molecular diffusion coefficient is consistent with the dispersivity defined as the fraction of surrounding molecules that moved away after 1 μs of simulation (Fig. S10a). This parameter may also serve as an indicator of the fusion speed, as fusion, similar to dispersion, involves breaking and reforming intermolecular contacts. The dispersivities have the following order: F < L < M < P, which is precisely the order of the measured fusion speeds (Fig. 4f).

## Discussion

By combining brightfield, confocal, and negative-staining electron microscopy, optical tweezers, and all-atom molecular dynamics simulations, we have characterized the condensates formed by homo-tetrapeptides of eight nonpolar amino acids. The condensates have a variety of morphologies, including droplets, amorphous aggregates, gels, and amorphous dense liquids; the latter appear to have not been reported previously. The phase-separation threshold concentrations of these peptides differ by 368-fold, from 0.25 mg/mL for WWssWW to 92 mg/mL for PPssPP. The droplets formed by three of the tetrapeptides span a wide range of material properties, including a 736-fold difference in fusion speed and a 3856-fold difference in viscosity.

For these homo-tetrapeptides of nonpolar amino acids, we have found that the size of amino acid, by serving as a measure of sidechain interaction strength, is a strong determinant of the phase-separation threshold concentration, fusion speed, and viscosity. An important advantage of the short peptides is that their small sizes enable all-atom molecular dynamics simulations of spontaneous phase separation and two-phase equilibrium and the calculation of equilibrium and dynamic properties in the separated phases. The calculated phase-separation threshold concentrations are comparable to the experimental counterparts, and the differences in calculated dispersivities and diffusion coefficients qualitatively explain the observed disparities in fusion speed and viscosity. Expanding the present approach to all the 20 natural amino acids, to hetero-tetrapeptides and tetrapeptide mixtures may ultimately allow us to precisely determine the contributions of individual amino acids to phase equilibrium and material properties of IDP condensates.

The sidechain interaction strength, as indicated by the size of amino acid, is also a contributing factor to condensate morphology. Specifically, the tendency to form three of the condensate morphologies: amorphous aggregate, droplet, and ADL, appears to be correlated with interaction strength. Strong interactions (as in WWssWW) favor amorphous aggregates, medium interactions (as in FFssFF, LLssLL, and MMssMM) favor droplets, and weak interactions (as in

AAssAA and PPssPP) favor ADLs. However, the fourth morphology observed here, gel, cannot be solely explained by interaction strength. Backbone hydrogen bonding may possibly be an additional determinant for condensate morphology, in particular for gel formation. Our study raises a number of questions about gels. Why are amorphous aggregates self-limiting in size but gels grow indefinitely? What are the molecular structures that underlie the filamentous features of gels?

Our study also highlights the importance of criticality in biomolecular condensates, which has received only scant attention ^20, 34, 41^. We have shown that five of the tetrapeptides can form amorphous dense liquids, suggesting that this morphology may occur widely for biomolecular condensates. Several lines of evidence indicate that ADLs are metastable precursors to liquid droplets, due to critical slowing down. We hypothesize that some biomolecular condensates, e.g., transcription condensates ^42^, may include ADLs in their mix.

More generally, ADLs provide a condensed form distinct from liquid droplets for cellular functions. Lastly, the fact that fusion speeds exhibit critical scaling behavior means that environmental factors such as pH may sensitively tune material properties.

## Methods

### Synthesis, purification, and characterization

The procedure for the tetrapeptides with X = F, M, L, I, V, and W followed Abbas et al. ^17^; modifications were introduced for X = A and P. Details are described in Supplementary Methods, including Figs. S14-S15 for ^1^H and ^13^C NMR and mass spectra. The procedure for the dye molecule, 2-[4-(diethylamino) styryl]-3,3-dimethyl-1-octadecyl-3H-indol-1-ium iodide, was as described by Wang et al. ^43^. Details are found in Supplementary Methods, including Fig. S16 for ^1^H and ^13^C NMR spectra.

### Preparation of tetrapeptide condensate samples

The stock solution of a known concentration was prepared by weighing the white power of a peptide on an analytical balance and dissolving in milli Q water or 50 mM imidazole buffer. A few drops of 37% (w/w) HCl were added to lower the initial pH to 2. The stock solution was filtered with a Millex polyethersulfone syringe filter (0.22 μm pore size; Millipore Sigma catalog # SLGPR33RS) and stored at room temperature. An aliquot was diluted to the desired concentration (typically 20 μL final volume) with either milli Q water or the buffer; condensate formation was induced by raising pH with the addition of a small amount of 5 M NaOH.

### Brightfield microscopy

A 2-μL drop was placed on a glass slide and observed under an Olympus BX53 brightfield microscope using a 40× objective. Images were exported in tiff format and processed using ImageJ. For studying the effects of urea, condensate samples were prepared with urea added before raising pH.

### Negative-staining electron microscopy

A 20-μL drop of tetrapeptide sample, a 20-μL drop of 1% phosphotungstic acid (1%; Electron Microscopy Sciences SKU 19502-1), and three 20-μL drops of deionized water were sequentially placed on a piece of parafilm. Using forceps, a formvar-coated grid (Ted Pella, product # 01753-F) was positioned onto the sample drop for 2 min. Excess liquid was removed by gently touching the edge of the forceps with the ragged edge of a piece of filter paper. The grid was transferred to the phosphotungstic acid drop for 2 min, and then rinsed in the deionized water drops for 10 sec each. Surplus liquid was again removed by filter paper. The grid was left to dry for 10 min in a covered petri dish. EM images were acquired on a JEM-1400 Flash transmission electron microscope (JEOL) equipped with a BIOSPRINT 12M-B camera (AMT) operating at 80 kV.

### Confocal microscopy

All other experiments were conducted at room temperature on a LUMICKS C-Trap instrument, which includes a brightfield camera, a confocal fluorescence imaging module, and dual-trap optical tweezers. Confocal scanning over a 43.9 μm × 33.5 μm field of view was carried out with excitation wavelength at 532 nm (20% laser power; 100 nm pixel size; 0.0128 or 0.1 ms pixel dwell time). A brightfield image of the scanned area was also taken for comparison.

For FRAP experiments, a 200 nm × 200 nm square region inside condensates was bleached for 5 or 10 sec with 100% laser power. Fluorescence recovery was then monitored with 20% laser power and 0.05 ms pixel dwell time. The scanned area was 14.7 μm × 13.8 μm. After shifting and normalization by the pre- and post-bleaching fluorescence intensities, the recovery curve was fit to

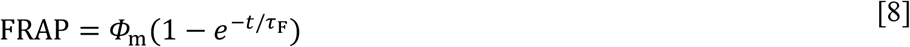

where *Φ*_m,_ denotes the mobile fraction and *τ*_F_ denotes the recovery time. When spontaneous photobleaching occurred, the decrease in fluorescence in a region away from the laser-bleached region was fit to a linear function of time and FRAP was further normalized by this function to correct for spontaneous photobleaching.

### OT-directed droplet fusion

Fusion speed was measured according to Ghosh and Zhou ^23^. Two droplets of equal size were trapped using 2% to 10% overall power and approximately 50:50 split. The trap-1 droplet was moved toward the trap-2 droplet at 10 nm steps until the droplets came into contact. The trapping forces were recorded at a sampling rate of 78125 Hz. The entire process was recorded by the brightfield camera; the video was analyzed using ImageJ to determine the radii of the droplets before and after fusion. The trap-2 force trace was fit to

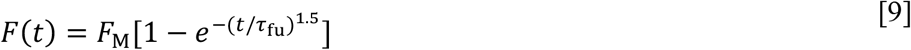

where *F*_M_ denotes the maximum force and *τ*_fu_ is the fusion time.

### Interfacial tension measurement by droplet stretching

This measurement followed Zhou ^35^ and Ghosh et al ^24^. Droplet samples containing uncoated polystyrene beads (1.13-μm radius) were prepared. Two droplets, each containing a single bead, were trapped and fused to generate a configuration where a single droplet was suspended by two trapped beads at the opposite poles. The overall trapping power was 10% to 40% and split 50:50 between the two traps. While fixing trap 2, trap 1 was pulled at a low speed of 0.05 μm/s for a distance of 0.5 μm. The pulling process was recorded by the brightfield camera; the video was analyzed using ImageJ to obtain the radius of the unstretched droplet and to verify that droplet stretching proceeded without interference. The spring constant of the droplet-trap system was obtained from the ratios of the trapping forces over the extension of the droplet. Using the stiffnesses of the two traps to remove their contributions, the spring constant, *χ*_0_, of the droplet was calculated. Lastly the interfacial tension of the droplet was found as

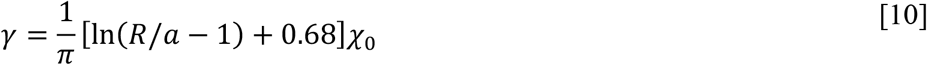

where *R* is the radius of the unstretched droplet and *a* is the radius of the beads.

### Condensate viscosity measurement by bead tracking

This measurement followed Kota and Zhou ^36^. Large droplets (> 5 μm in radius), each containing a single bead (1.13-μm radius) and settled on a coverslip coated with polyethylene glycol ^44^, were used for bead tracking. The bead was trapped and moved to the center of the droplet and then released. After waiting for 5-10 min, a brightfield image of the bead was taken to serve as the template, and the bead movement was then tracked by template matching at a 15 Hz frame rate. The time trace of the bead *x* and *y* positions was exported. The MSD was calculated and fit to

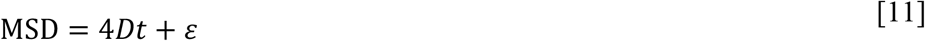

where *D* denotes the 2-dimensional diffusion constant of the bead and *ε* is a constant that arises from the uncertainty in locating the bead position ^45^. The zero-shear viscosity was then obtained according to the Stokes-Einstein relation

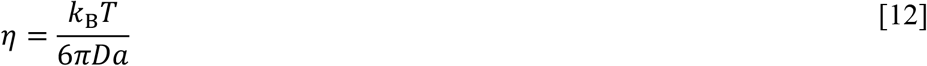

where *k*_3_ is the Boltzmann constant, *T* is the absolute temperature, and *a* again is the bead radius.

### Molecular dynamics simulations

The tetrapeptides, denoted as XXssXX, consist of two dipeptides linked by NH-CH_2_-CH_2_-S-S-CH_2_-CH_2_-NH (Fig. S1). We generated force-field parameters for the linker using Amber tools ^46^ and Gaussian 16 ^47^. More specifically, we parameterized one half of the linker and connected the two halves by a disulfide bond. The half linker patched with terminal hydrogens, NH_2_-CH_2_-CH_2_-SH, was generated using Pymol (https://pymol.org); its initial coordinate file for parametrization was prepared using Antechamber. Geometry optimization and partial charge calculation were performed at the HF/6-31G* level in Gaussian 16. Other parameters were from the Parm10 parameter file. prepgen was used to prepare the half linker as an amino-acid-like residue and connected it to a dipeptide to generate the reduced form, XXs-H, of the tetrapeptide. Any missing parameters were completed by parmchk. Lastly, a disulfide bond between two copies of the reduced form was imposed in tleap to form a tetrapeptide. The parameters for the link residues were appended to Amberff14SB ^48^ to form the force field of the tetrapeptides; parameters for neutral terminal residues were modified from the charged versions. The water model was TIP4PD ^49^.

Preparation of a dense solution of the tetrapeptides and subsequent simulations were done in AMBER18 ^50^ as described previously ^37^. Energy minimization (2000 steps of steepest descent and 3000 steps of conjugate gradient) was carried out using sander; All MD simulations were carried out on GPUs using *pmemd*.*cuda* ^51^. Long-range electrostatic interactions were treated using the particle mesh Ewald method ^52^ with a nonbonded cutoff of 10 Å. The Langevin thermostat with a damping constant of 3 ps^-1^ was used to maintain temperature, while the Berendsen barostat ^53^ was used to regulate the pressure. All bonds connected with the hydrogen atoms were constrained using the SHAKE algorithm ^54^.

To start, 8 copies of the tetrapeptide with both termini neutral (to model high pH) were randomly inserted into a cubic box with a side length of 30 Å and solvated with 500-600 waters. After energy minimization, a short simulation of 100 ps was performed at constant NVT and a timestep of 1 fs, with temperature ramping from 0 to 294 K over the first 40 ps and maintained at 294 K for the remaining 60 ps. The 8 copies of the peptide in the last snapshot were duplicated in each of three orthogonal directions to build 64 copies in a cubic box with a side length of 60 Å, solvated again with 3800-5100 water molecules. To condense the system, different levels of water molecules (1/4, 1/3, 1/2, and 2/3) were randomly removed, which generated four independent systems. These systems were energy minimized again and first simulated for 500 ps at constant NVT (temperature ramped from 0 to 294 K for 40 ps and then maintained at 294 K for 460 ps; 1 fs timestep). The simulations then continued in duplicates at constant NPT (294 K and 1 bar) for 3.6-6.8 μs at a timestep of 2 fs. The cubic simulation boxes settled to ∼54 Å in side length for the systems with 1/4 water removal to ∼47 Å with 2/3 water removal.

Slab configurations from the preceding simulations of FFssFF and LLssLL were modified to lower pH values and simulated in an elongated box (*L*_*z*_/*L*_*x*_ = 5). The p*K*_a_ values of the terminal amines were assumed to 7 and 6, respectively; the lower second p*K*_a_ models an increased tendency to be neutral once the first terminus is already charged. For pH 7.7, 18 copies of the peptide had one terminal amine changed from neutral to charged; this number increased to 32 copies at pH 7.3. For pH 6.4, 48 copies had one charged amine and 10 copies had two charged amines. The 64 copies of the peptides were solvated with 18400-22000 water molecules and neutralized with Cl^−^ ions. Each system was energy minimized and simulated first at constant NVT (500 ps at 1 fs timestep) and then at constant NPT (100 ns at 2 fs timestep). Lastly, the simulations continued at constant NVT for 4.5-4.9 μs each at a 2 fs timestep. Snapshots were saved at 100 ps intervals for analysis.

### Simulation data analyses

Backbone hydrogen bonds and the number of neighboring or surrounding chains (3.5 or 4 Å cutoff between heavy atoms) were calculated using CPPTRAJ ^55^ on a 1000-ns portion of the cubic-box simulations. This portion was chosen based on well-formed slabs for peptides that did form slabs. The calculations were done by centering the periodic system on each copy of the peptide and then averaging over all the 64 copies.

For the elongated-box simulations, binodals were calculated using tcl scripts and MSDs were calculated using CPPTRAJ and python scripts. For binodals, the simulation box was divided into 2-Å thick slices along the *z* axis and the number of peptide heavy atoms in each slice was counted. The results were averaged over the snapshots in the 1000-4500 ns portion of the simulations. The heavy-atom count as a function of the *z* coordinate was fit to a hyperbolic tangent function to obtain the densities in the dense and dilute phases. MSDs were calculated over the 3490-4530 ns portion of the simulations at pH 7.3. For the dilute phase, calculations were done on the unwrapped trajectories of the peptide molecules that stayed in the dilute phase over the considered time period. For the dense phase, after unwrapping, the center of the chains that remained in the slab was removed before calculating MSDs. VMD ^56^ and ChimeraX ^57^ were used for rendering images and making movies.

## Acknowledgments

We thank Divya Kota for assistance with confocal microscopy, Fidha Muhammedkutty for assistance with simulation data analysis, and Andy Nguyen for the use of an HPLC instrument.

## Author contributions

Conceptualization: HXZ, YZ, RP

Methodology: YZ, RP, SS, DL

Investigation: YZ, RP, HXZ

Funding acquisition: HXZ

Project administration: HXZ

Supervision: HXZ

Writing: HXZ, YZ, RP, SS

## Supplementary information

Supplementary Figures S1 to S13

Supplementary Methods, including Figures S14 to S16

Supplementary Movie S1 to S3

