## Supplementary Figures and Methods for "Amino Acid-Dependent Material Properties of Tetrapeptide Condensates"

*Supplementary Materials*

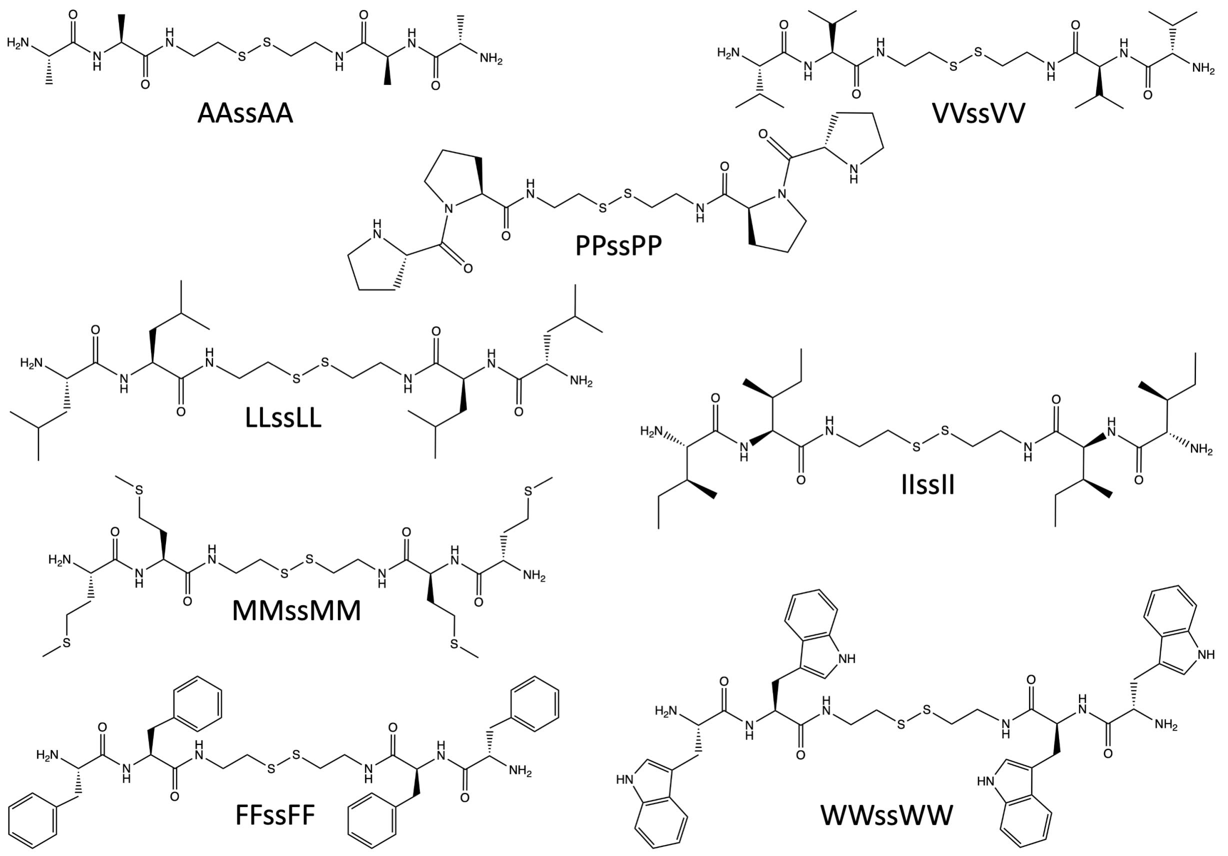


Fig. S1 Chemical structures of the eight tetrapeptides.

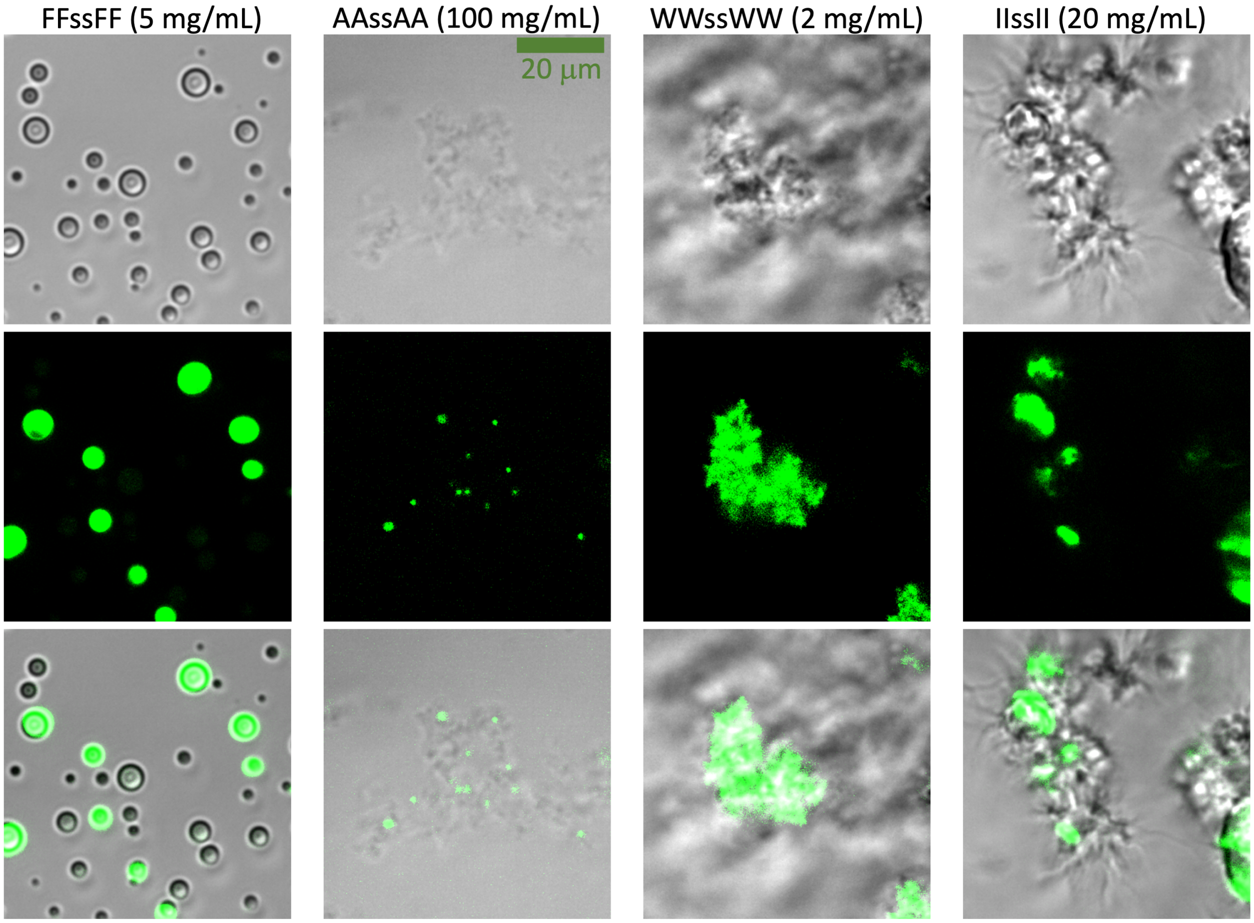


Fig. S2 Confocal images of tetrapeptide condensates formed at the indicated concentrations in milli Q water and pH 13. The dye concentration was 10 µM. First row: brightfield images; second row: confocal images; third row: merge. The pixel dwell time was 0.0128 ms for three of the peptides and increased to 0.1 ms for WWssWW.

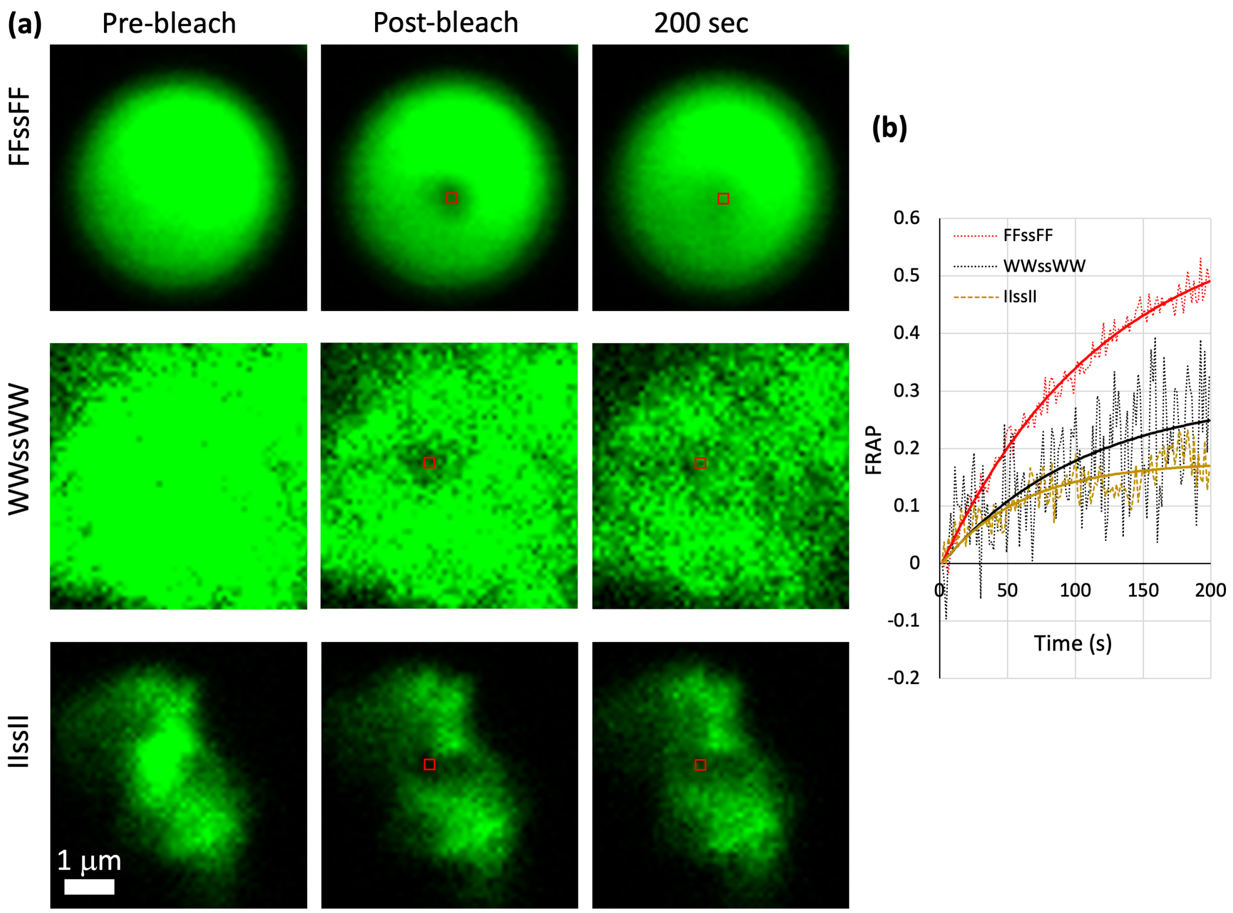


Fig. S3 FRAP results for three tetrapeptide condensates. (a) Confocal images prior to photobleaching, immediately after photobleaching, and after 200 sec of recovery. A red square indicates the bleached region. Samples were prepared as described in Fig. S2. The photobleaching time was 5 sec for FFssFF and WWssWW and 10 sec for IIssII. (b) Recovery curves, shifted and normalized by the pre- and post-bleach intensities. Correction for spontaneous photobleaching was applied to WWssWW and IIssII. The large fluctuations in the WWssWW curve are due to movement of the aggregate during confocal scanning. Dotted curves show raw data; solid curves show fits to Eq [8].

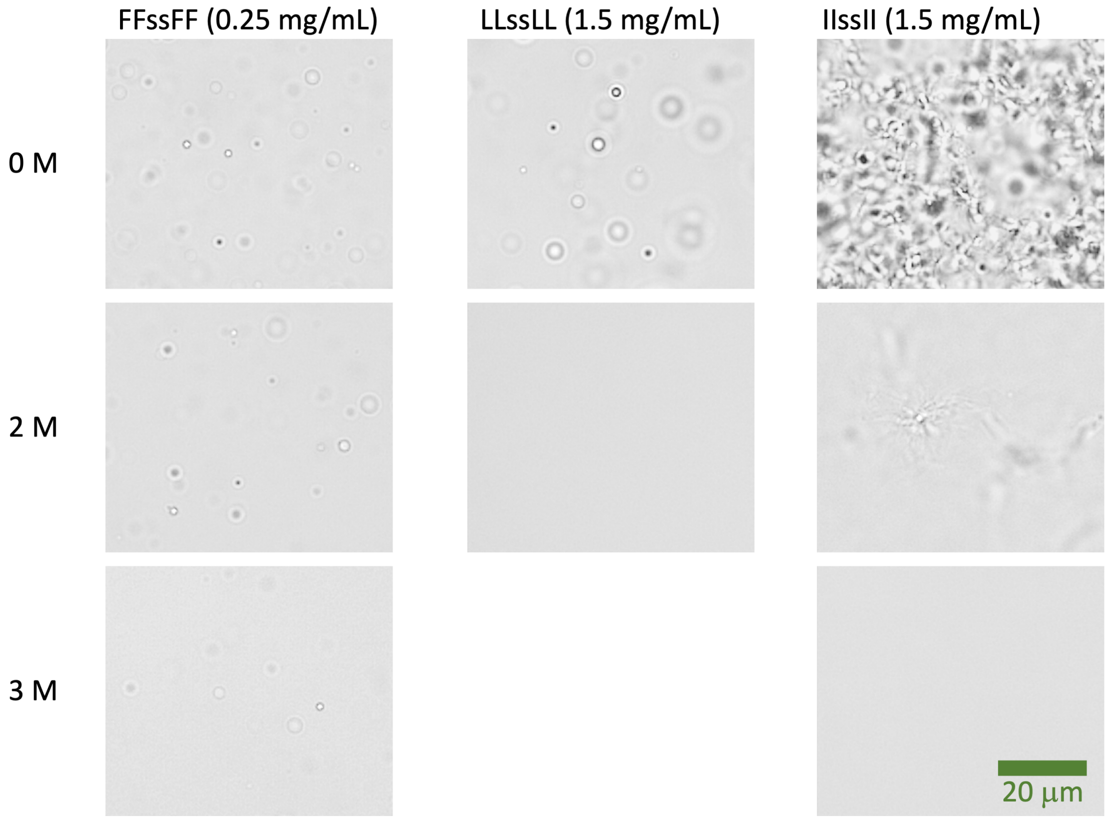


Fig. S4 Effects of urea on the phase separation of three tetrapeptides. Condensates were formed at the indicated peptide and urea concentrations in milli Q water and pH 13. FFssFF and LLssLL samples were observed immediately after raising the pH to 13; for IIssII, gels took 5 min to emerge at 2 M urea and were never observed at 3 M urea.

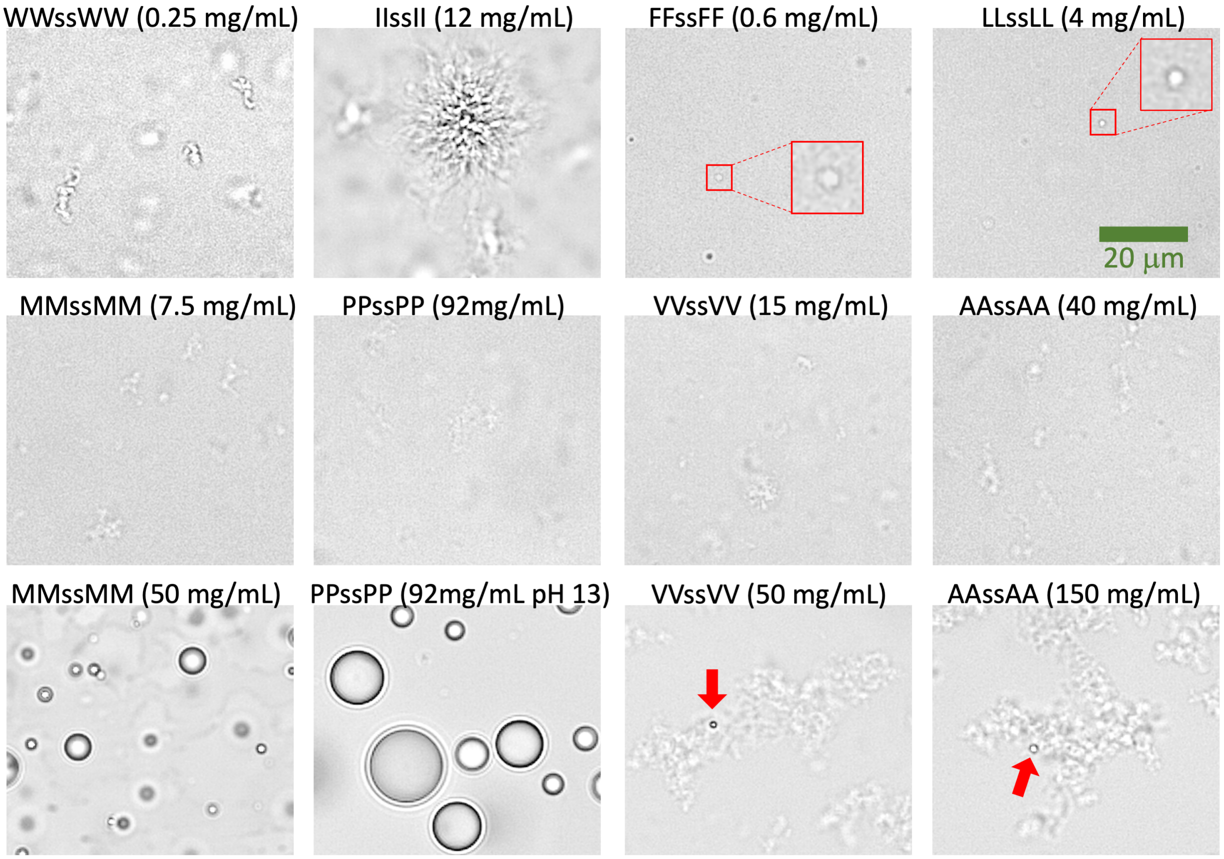


Fig. S5 Brightfield images showing the condensate morphologies at the threshold concentrations of eight tetrapeptides and pH 7 in 50 mM imidazole buffer. For the first two rows, samples of seven peptides were observed immediately after raising the pH; the IIssII sample was observed after 5 min of incubation. In the last row, the concentration or pH is raised, showing either the conversion from ADLs to droplets (MMssMM and PPssPP) or an increased tendency to form droplets inside ADLs (VVssVV and AAssAA).

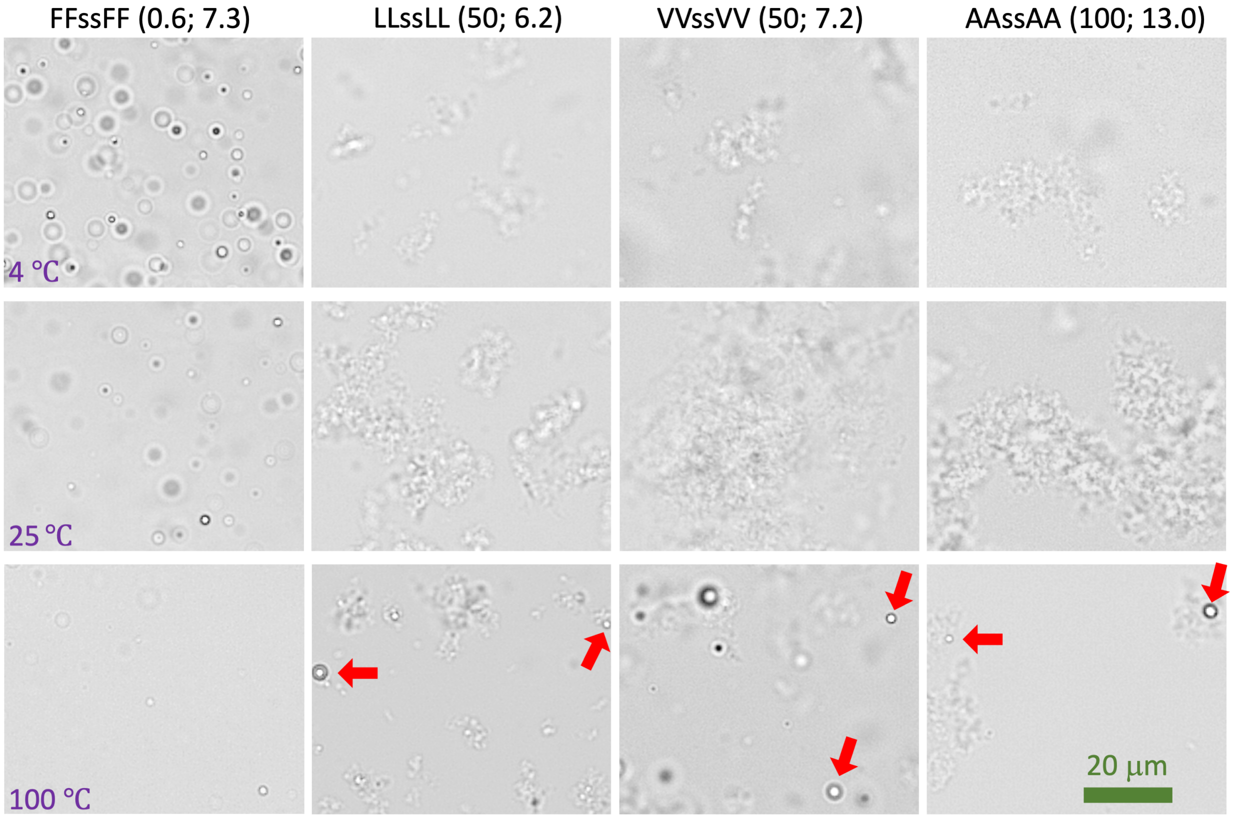


Fig. S6 Effects of temperature on phase separation and condensate morphology. Condensates were formed at the indicated tetrapeptide concentrations [first number (in mg/mL) inside parentheses] and pH (second number inside parentheses). In the last row, droplets are indicated by red arrows. For 4 °C, 60 μL of sample was placed in ice for 2-3 min before raising pH; a 2-μL aliquot was then immediately observed under a microscope. The protocol for 100 °C was similar, except the sample was placed in a heating block.

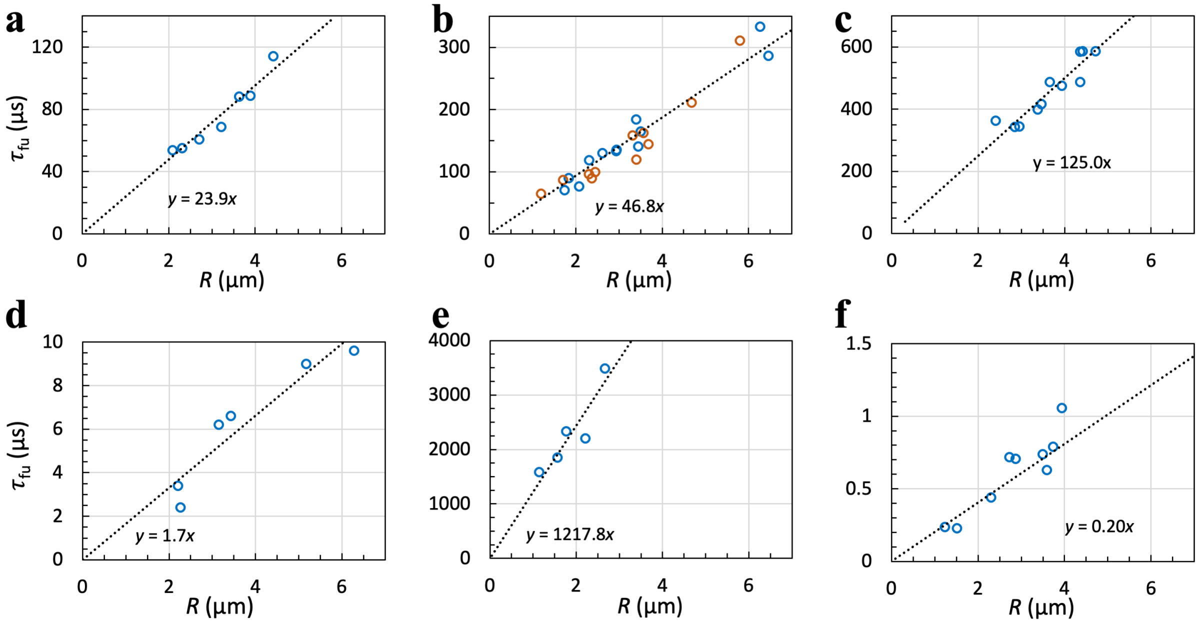


Fig. S7 Fusion speeds of tetrapeptide droplets. (a) LLssLL at 5.5 mg/mL and pH 12. (b) LLssLL at 5.5 (brown circles) or 30 mg/mL (blue circles) and pH 8. (c) LLssLL at 30 mg/mL and pH 7. (d) MMssMM at 50 mg/mL and pH 10. (e) FFssFF at 4 mg/mL and pH 10. (f) PPssPP at 167 mg/mL and pH 13.

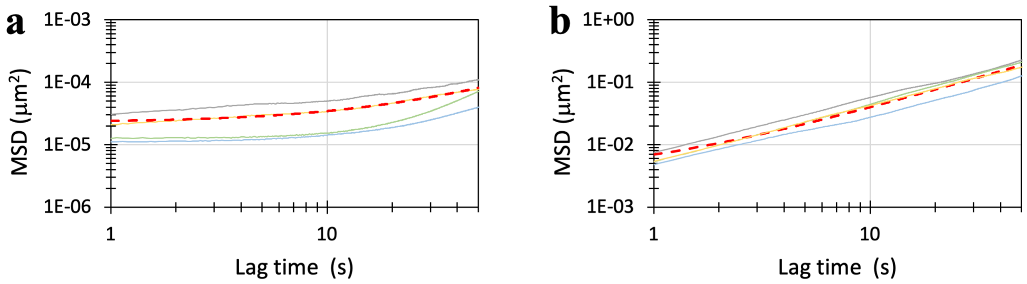


Fig. S8 Mean-square-displacements of a bead inside tetrapeptide droplets. (a) FFssFF at 4 mg/mL and pH 10. (b) MMssMM at 50 mg/mL and pH 10.

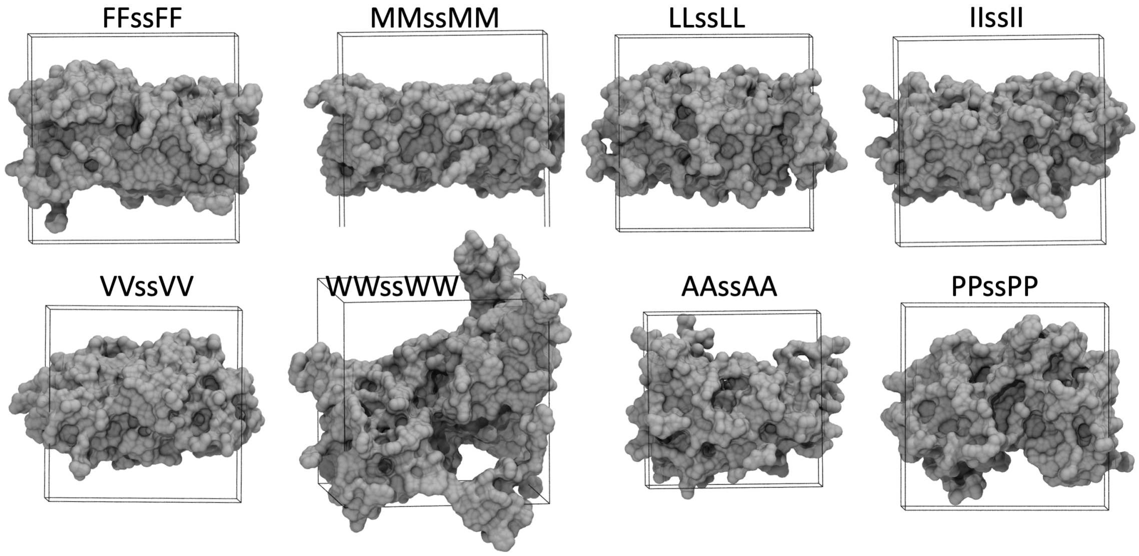


Fig. S9 Condensate morphologies in molecular dynamics simulations. The snapshots are the same as those in Fig. 7a, but are rendered using molecular surface.

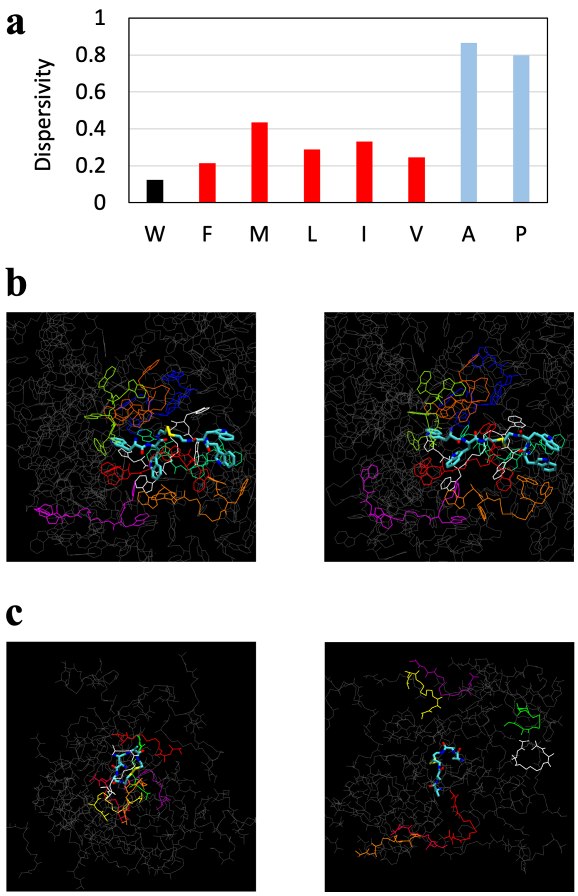


Fig. S10 Dispersivities of tetrapeptides in molecular dynamics simulations. (a) Comparison of dispersivities among eight peptides. (b) Snapshots at the start (left) and end (right) of a 1-µs portion of a simulation for WWssWW. A tagged peptide molecule is rendered as stick; its surrounding molecules (with a 4-Å cutoff between heavy atoms) at the start are rendered as line. These snapshots are the same as the first and last frames of Movie S2. (c) Corresponding results for AAssAA. The snapshots are the same as the first and last frames of Movie S3.

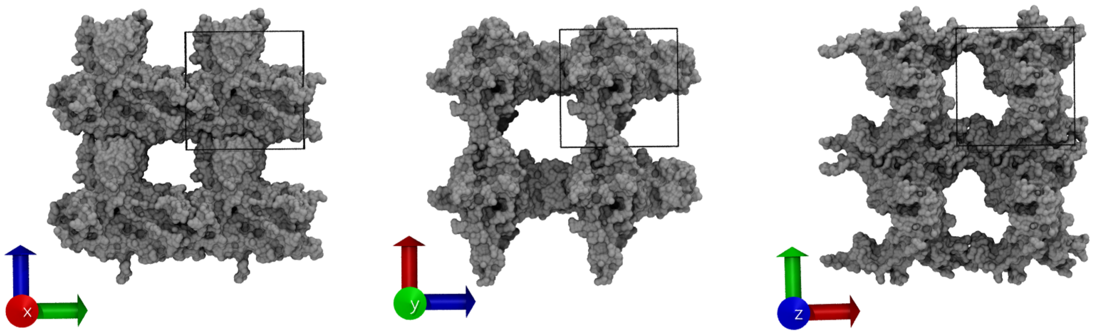


Fig. S11 Gel-like condensates in molecular dynamics simulations of IIssII. For each of the three orthogonal viewing directions, a unit cell (indicated by a box) plus three image cells are shown; a hole is formed in each direction. Two other gel-like forms were formed in six other simulations: in one, holes are formed in two of the three directions; in the other, a slab with a hole is formed.

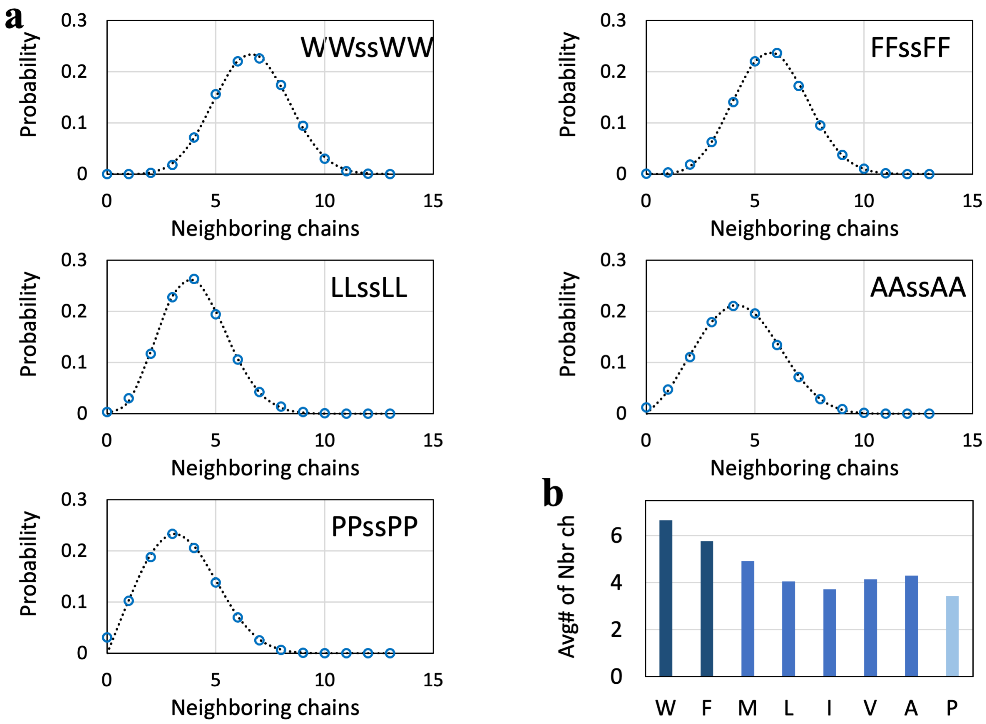


Fig. S12 Distribution of the number of neighboring chains. (a) Distributions for five tetrapeptides. Circles are raw data from the simulations; curves are fits to the generalized Gamma distribution. (b) Averages values of the number of neighboring chains. Chain neighbors were defined with a 3.5-Å cutoff between heavy atoms.

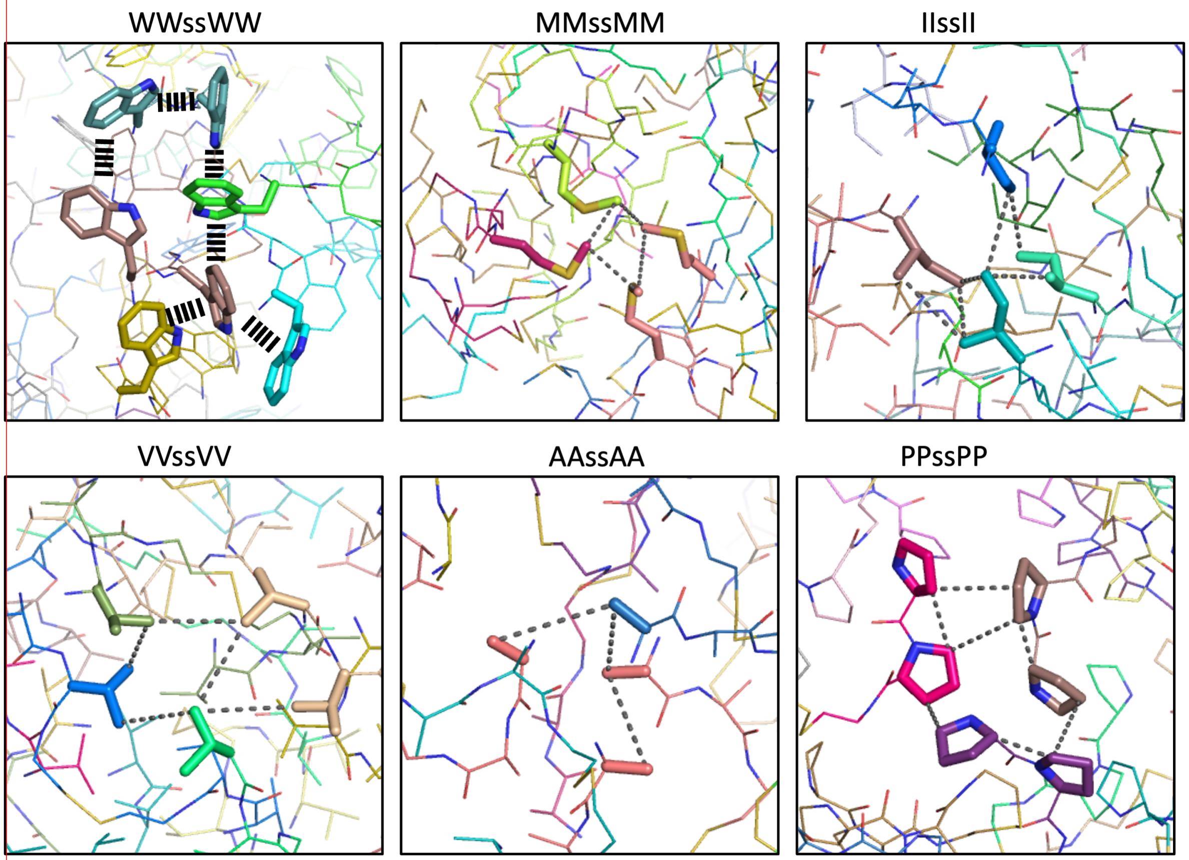


Fig. S13 Intermolecular interactions in molecular dynamics simulations of tetrapeptides.

### Supplementary Methods

*1. Peptide synthesis, purification, and characterization*

**Materials.** Boc-L-alanine (Boc-A-OH), Boc-L-proline (Boc-P-OH), Boc-L-valine (Boc-V-OH), Boc-L-leucine (Boc-L-OH), Boc-L- isoleucine (Boc-I-OH), Boc-L-methionine (Boc-M-OH), Boc-L-phenylalanine (Boc-F-OH), Boc-L- tryptophan (Boc-W-OH), 1-hydroxybenzotriazole (HOBt), *N,N,N′,N′*-tetramethyl-*O*-(1*H*-benzotriazol-1-yl)uronium hexafluorophosphate (HBTU), *N*,*N*-diisopropylethylamine (DIPEA), cystamine dihydrochloride (CDC), hydrogen chloride solution (4 M in dioxane), and all the solvents used were from Sigma Aldrich.

**Synthesis.** (1) Amide coupling. Boc-X-OH (3.1 mmol, 1 equiv.), HOBt (398 mg, 2.94 mmol, 0.95 equiv.), HBTU (1114 mg, 2.94 mmol, 0.95 equiv.), and 5 mL of dimethylformamide (DMF) were mixed in a round-bottom flask. Then, DIPEA (2.1 mL, 12.4 mmol, 4 equiv.) and CDC (316 mg, 1.4 mmol, 0.45 equiv.) were added and the reaction mixture was stirred at room temperature until completion [typically 24 hours, monitored by thin layer chromatography (TLC)). For X = F, M, L, I, V, and W, ~80 mL of water was added to the mixture, and centrifugation (in two 50-mL tubes at 4500 RPM for 30 minutes) was used to collect the white precipitate. For X = A, the reaction mixture was dissolved in diethyl ether; Boc-AssA-Boc was separated by washing with water three times and concentrated in a rotary evaporator; further separation was performed by column chromatography ($CH_2Cl_2$/MeOH, 15:1 ratio). For X = P, separation was done by preparative high-performance liquid chromatography (HPLC) (see below). (2) Hydrolysis. The material (Boc-XssX-Boc) from the previous step was dissolved in dioxane/dichloromethane (DCM) (3 mL, 4:1 ratio) in a round-bottom flask. Hydrogen chloride (4 M in dioxane, 6 mL) was then added, and the reaction was kept at room temperature for 3 hours. For X = F, M, L, I, V, W, and A, the reaction mixture was then concentrated in a rotary evaporator. The resulting yellow oil was dissolved in 70 mL of diethyl ether, and the white precipitate was collected after centrifugation. For X = P, separation was again done by preparative HPLC. (3) Formation of XXssXX. Step (1) was performed again, with the material (XssX) from step (2) replacing CDC; this time only the separation for X = P was the exception. Lastly, hydrolysis was carried out following the protocol step (2).

**Purification.** Final purification for the other seven peptides (other than X = P) was conducted by preparative HPLC using a Prep 2545 HPLC system (Waters), which was outfitted with a C18 column (XBridge Peptide BEH; Waters). The HPLC process was operated at a flow rate of 17 mL/min and was monitored at a wavelength of 214 nm. The final product was obtained by lyophilization.

**Characterization.** $^{1}H$ and $^{13}C$ NMR spectra were recorded on a Bruker AV-500 spectrometer. $^{1}H$ and $^{13}C$ NMR chemical shifts ($\delta$) are reported in parts per million (ppm) downfield of tetramethylsilane and referenced relative to, respectively, the residual solvent peak (2.50 ppm) and the carbon resonance (39.52 ppm) of the solvent [dimethyl sulfoxide-$d_6$ (DMSO-$d_6$)]. Multiplicities are indicated by s (singlet), brs (broad singlet), d (doublet), t (triplet), q (quartet), p (quintet), h (sextet), dd (doublet of doublet), dt (doublet of triplet), m (multiplet), and so on. $^{1}H$ NMR signals that fall within a ~0.3 ppm range are generally reported as a multiplet, with a range of chemical shift values corresponding to the peak or center of the peak. Coupling constants ($J$) are reported in Hz. Electrospray ionization (ESI) mass spectra were recorded on Micromass Q-Tof Ultima API (Waters) at University of Illinois Urbana-Champaign. Yields refer to the product before HPLC purification (except for X = P).

*NMR spectra of tetrapeptides*

a. AAssAA (410 mg, 67.1 %); white powder; **$^1$H NMR** (500 MHz, DMSO-d$_6$) $\delta$ 8.71 (d, $J$ = 7.5 Hz, 2H), 8.35 (d, $J$ = 5.8 Hz, 4H), 8.30 (t, $J$ = 5.7 Hz, 2H), 4.25 (p, $J$ = 7.1 Hz, 2H), 3.85 (p, $J$ = 6.2 Hz, 2H), 3.32 (dp, $J$ = 35.5, 6.8 Hz, 4H), 2.83 – 2.69 (m, 4H), 1.35 (d, $J$ = 7.0 Hz, 6H), 1.24 (d, $J$ = 7.1 Hz, 6H); **$^{13}$C NMR** (125 MHz, DMSO-d$_6$) $\delta$ 171.87, 169.08, 48.60, 48.09, 37.98, 37.17, 18.39, 17.16.

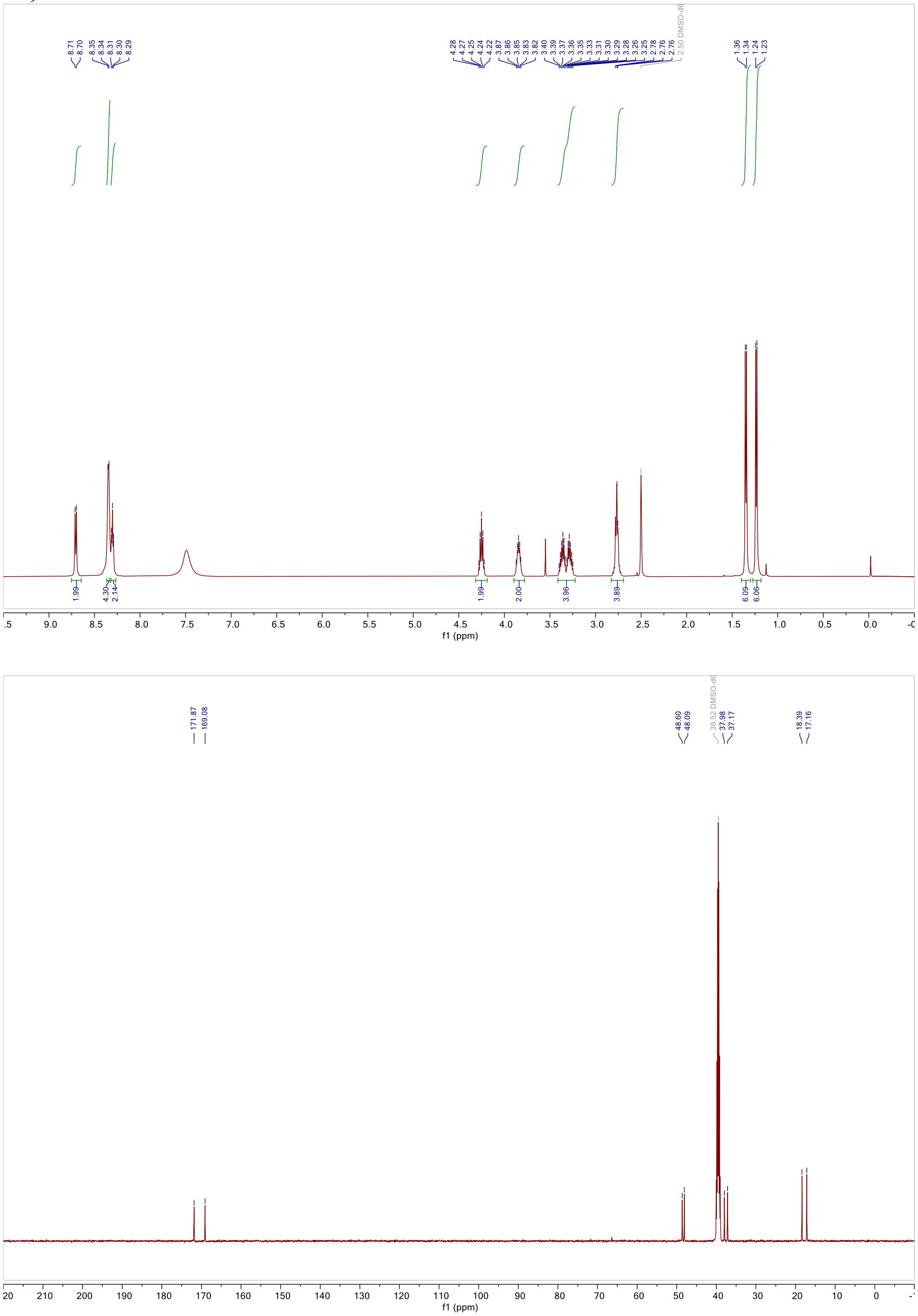


Fig. S14a $^1$H NMR (top) and $^{13}$C NMR (bottom) spectra of AAssAA in DMSO-d$_6$.

b. PPssPP (406 mg, 53.6 %); white powder; **$^1$H NMR** (500 MHz, DMSO-d$_6$) $\delta$ 8.54–8.41 (m, 2H), 8.16 (t, $J$ = 5.7 Hz, 2H), 4.47 (s, 2H), 4.31 (dd, $J$ = 8.4, 5.2 Hz, 2H), 3.63 (dt, $J$ = 10.0, 6.7

Hz, 2H), 3.47–3.43 (m, 4H), 3.39 (q, *J* = 6.7 Hz, 2H), 3.30–3.20 (m, 4H), 3.17 (s, 2H), 2.75 (td, *J* = 6.8, 4.1 Hz, 4H), 2.40 (dq, *J* = 12.7, 5.2 Hz, 2H), 2.13 (dq, *J* = 12.5, 7.3 Hz, 2H), 1.93–1.84 (m, 8H), 1.78 (dp, *J* = 12.4, 5.9 Hz, 2H); **$^{13}$C NMR** (125 MHz, DMSO-$d_6$) $\delta$ 171.65, 167.01, 60.33, 58.83, 47.21, 46.26, 38.27, 37.59, 29.82, 28.22, 24.89, 23.95.

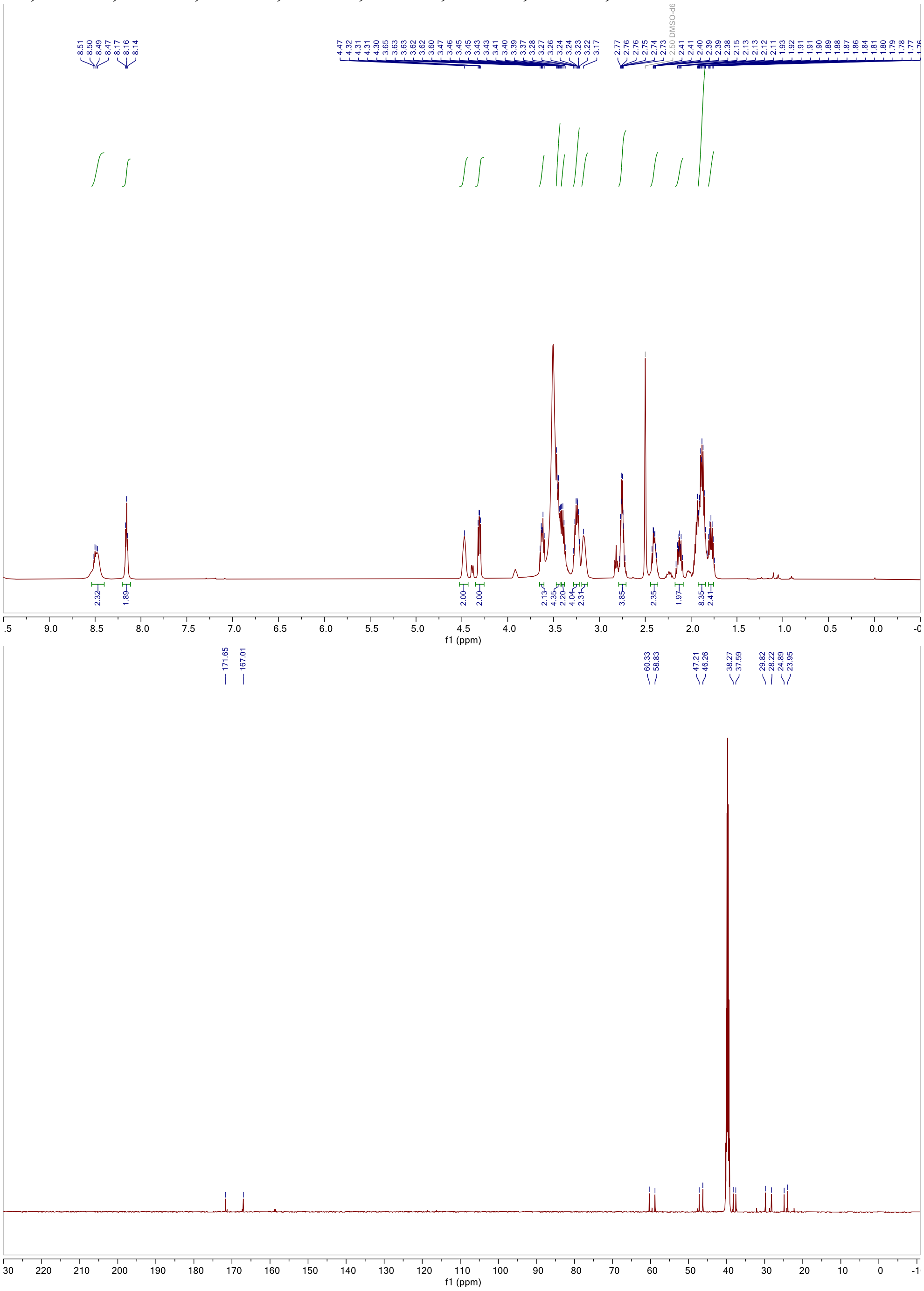

Fig. S14b $^1$H NMR (top) and $^{13}$C NMR (bottom) spectra of PPssPP in DMSO-$d_6$.

c. VVssVV (427 mg, 55.6 %); white powder; **$^1$H NMR** (500 MHz, DMSO-$d_6$) δ 8.49 (d, *J* = 8.6 Hz, 2H), 8.40 (t, *J* = 5.6 Hz, 2H), 8.30 (s, 4H), 4.12 (t, *J* = 7.8 Hz, 2H), 3.73 (t, *J* = 5.5 Hz, 2H), 3.36 (q, *J* = 6.6 Hz, 2H), 3.33–3.22 (m, 2H), 2.75 (t, *J* = 6.7 Hz, 4H), 2.07 (h, *J* = 6.7 Hz, 2H), 1.95 (h, *J* = 6.8 Hz, 2H), 0.90 (t, *J* = 5.6 Hz, 12H), 0.87 (t, *J* = 5.6 Hz, 12H); **$^{13}$C NMR** (125

MHz, DMSO-d$_6$) $\delta$ 170.99, 168.09, 58.70, 57.44, 38.23, 37.57, 30.97, 30.37, 19.67, 18.88, 18.73, 18.35.

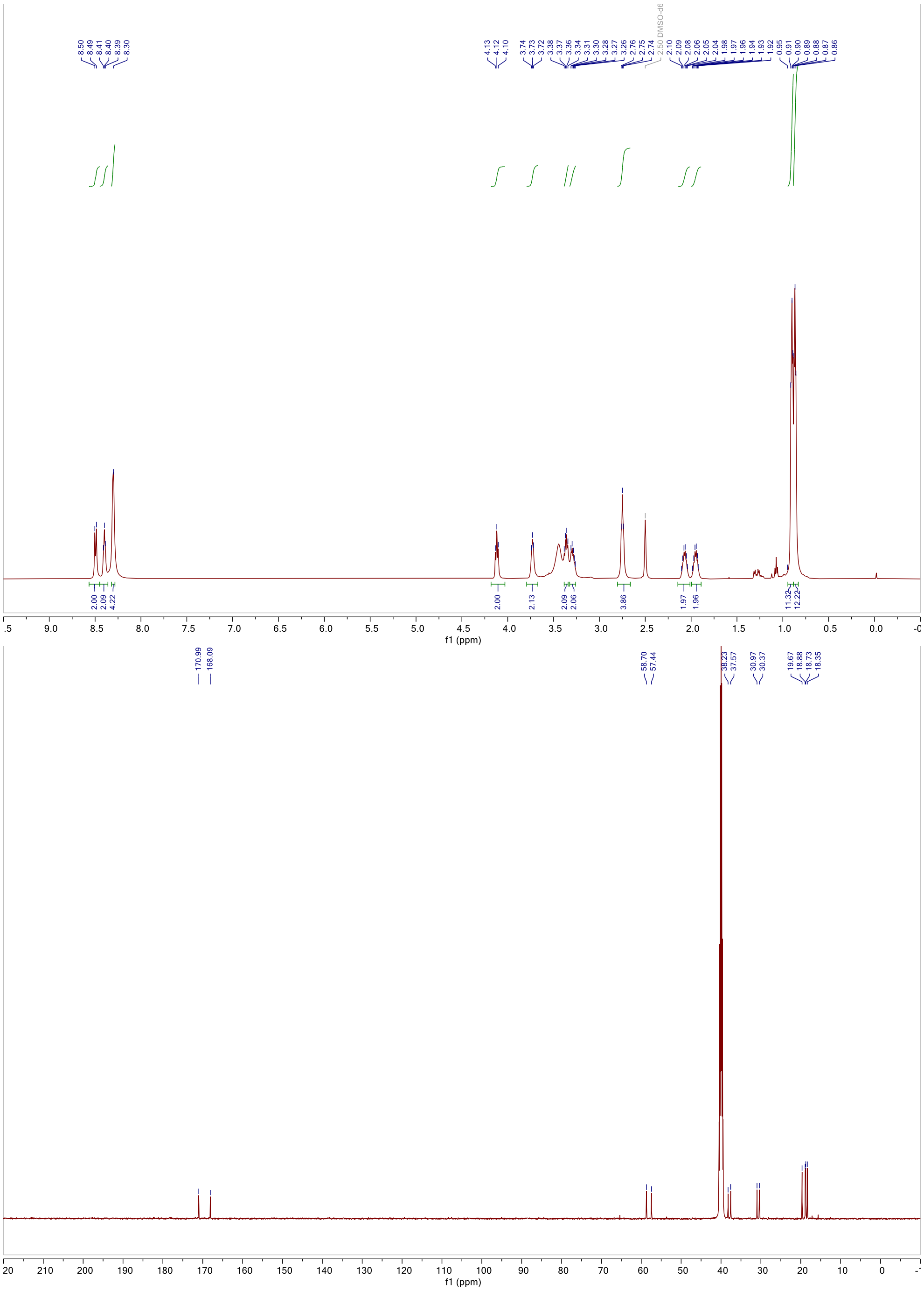


Fig. S14c $^{1}$H NMR (top) and $^{13}$C NMR (bottom) spectra of VVssVV in DMSO-d$_6$.

d. LLssLL (498 mg, 58.8 %); white powder; **$^{1}$H NMR** (500 MHz, DMSO-d$_6$) $\delta$ 8.57 (d, $J$ = 8.2 Hz, 2H), 8.29 (t, $J$ = 5.6 Hz, 2H), 8.09 (s, 4H), 4.34 (td, $J$ = 8.3, 6.3 Hz, 2H), 3.78 (s, 2H), 3.30 (dq, $J$ = 13.1, 6.5 Hz, 4H), 2.75 (td, $J$ = 6.6, 3.6 Hz, 4H), 1.61 (ddt, $J$ = 15.8, 12.3, 6.4 Hz, 4H), 1.52 (dq, $J$ = 13.7, 7.0 Hz, 4H), 1.48–1.41 (m, 4H), 0.92–0.82 (m, 24H); **$^{13}$C NMR** (125 MHz, DMSO-d$_6$) $\delta$ 171.86, 169.03, 51.62, 51.16, 41.71, 40.70, 38.23, 37.49, 24.50, 23.91, 23.33, 23.16, 22.33, 22.29.

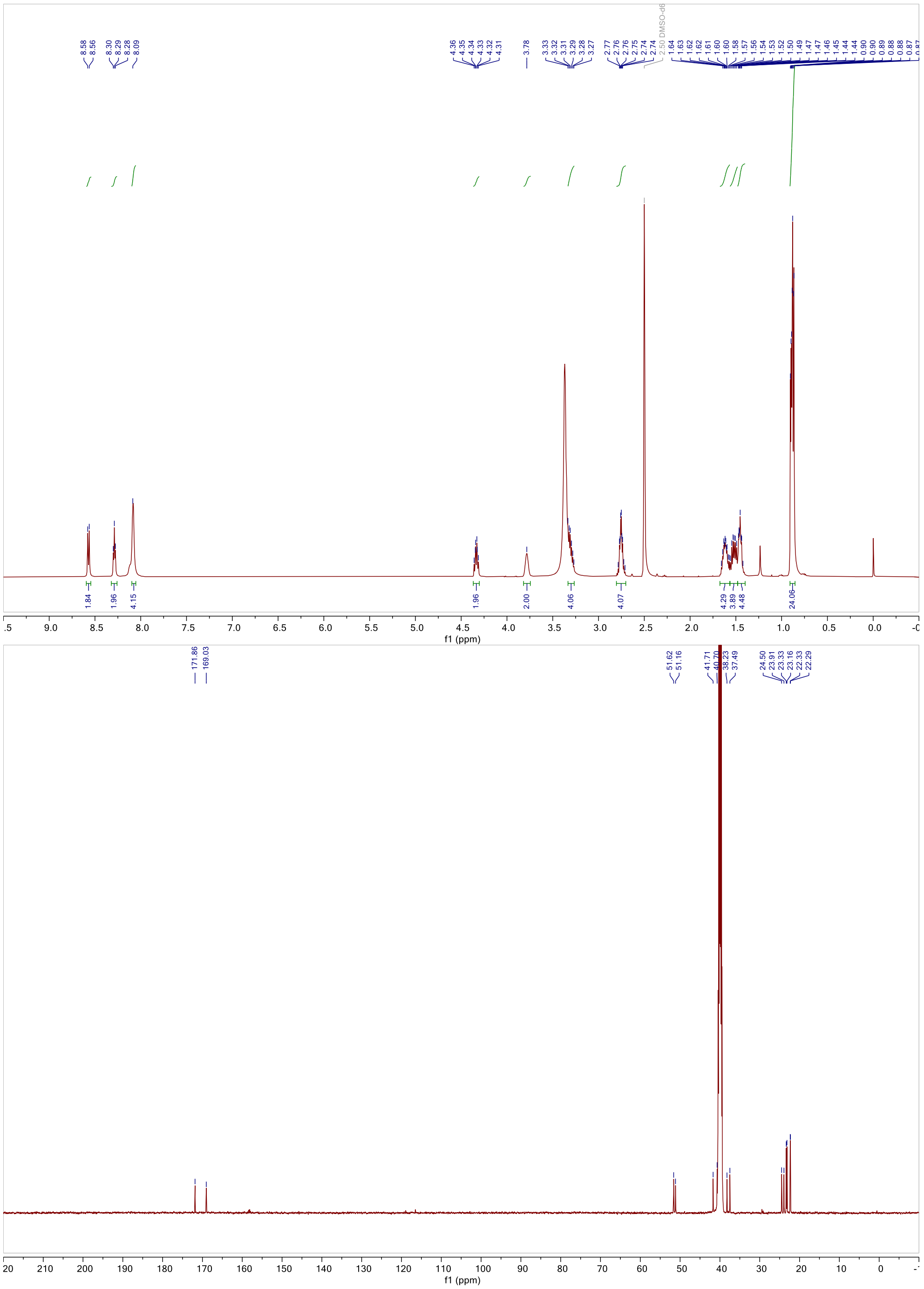


Fig. S14d $^{1}H$ NMR (top) and $^{13}C$ NMR (bottom) spectra of LLssLL in DMSO-$d_6$.

e. IIssII (511 mg, 60.3 %); white powder; **$^{1}$H NMR** (500 MHz, DMSO-$d_6$) $\delta$ 8.48 (d, $J$ = 9.1 Hz, 2H), 8.38 (t, $J$ = 5.3 Hz, 2H), 8.28 (s, 4H), 4.15 (t, $J$ = 8.2 Hz, 2H), 3.73 (s, 2H), 3.36–3.31 (m, 4H), 2.75 (t, $J$ = 6.6 Hz, 4H), 1.81 (s, 2H), 1.71 (s, 2H), 1.49 (s, 4H), 1.16–1.01 (m, 4H), 0.83 (m, 24H); **$^{13}$C NMR** (125 MHz, DMSO-$d_6$) $\delta$ 170.48, 167.46, 57.15, 56.04, 37.69, 37.00, 36.47, 36.17, 24.30, 23.93, 15.23, 14.35, 11.06, 10.97.

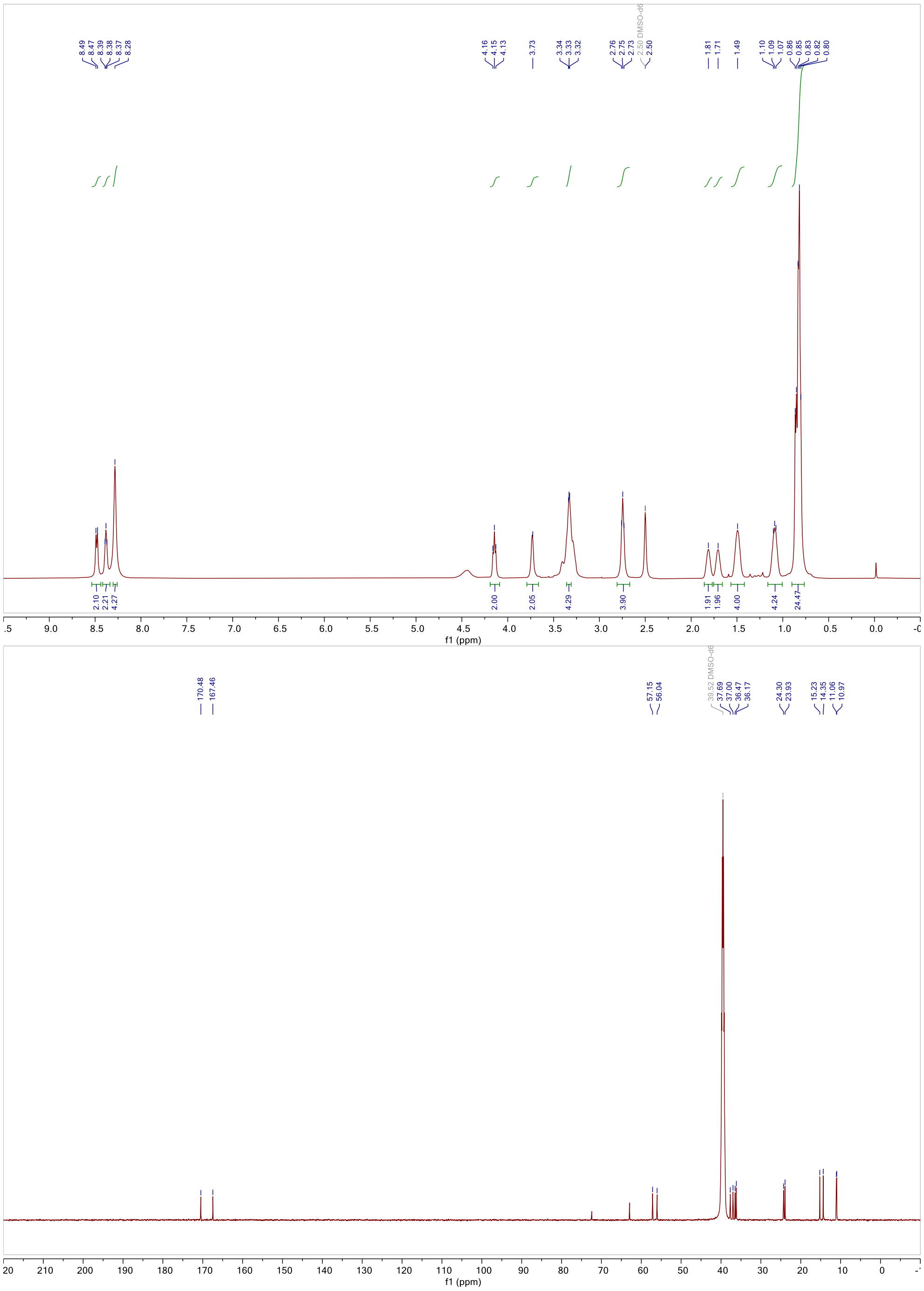


Fig. S14e $^{1}$H NMR (top) and $^{13}$C NMR (bottom) spectra of IIssII in DMSO-$d_6$.

f. MMssMM (632 mg, 66.7 %); white powder; **$^{1}$H NMR** (500 MHz, DMSO-$d_6$) $\delta$ 8.83 (d, $J$ = 7.7 Hz, 2H), 8.46 (s, 4H), 8.39 (t, $J$ = 5.2 Hz, 2H), 4.30 (td, $J$ = 8.3, 4.6 Hz, 2H), 3.93–3.86 (m, 2H), 3.57 (s, 4H), 3.37 (p, $J$ = 6.9 Hz, 2H), 3.27 (dt, $J$ = 13.4, 6.4 Hz, 2H), 2.76 (s, 4H), 2.47–2.39 (m, 2H), 2.02 (s, 12H), 1.98–1.86 (m, 8H), 1.86–1.76 (m, 2H); **$^{13}$C NMR** (125 MHz, DMSO-$d_6$) $\delta$ 171.05, 168.44, 52.56, 51.96, 38.26, 37.47, 32.37, 31.43, 29.90, 28.51, 15.04, 14.97.

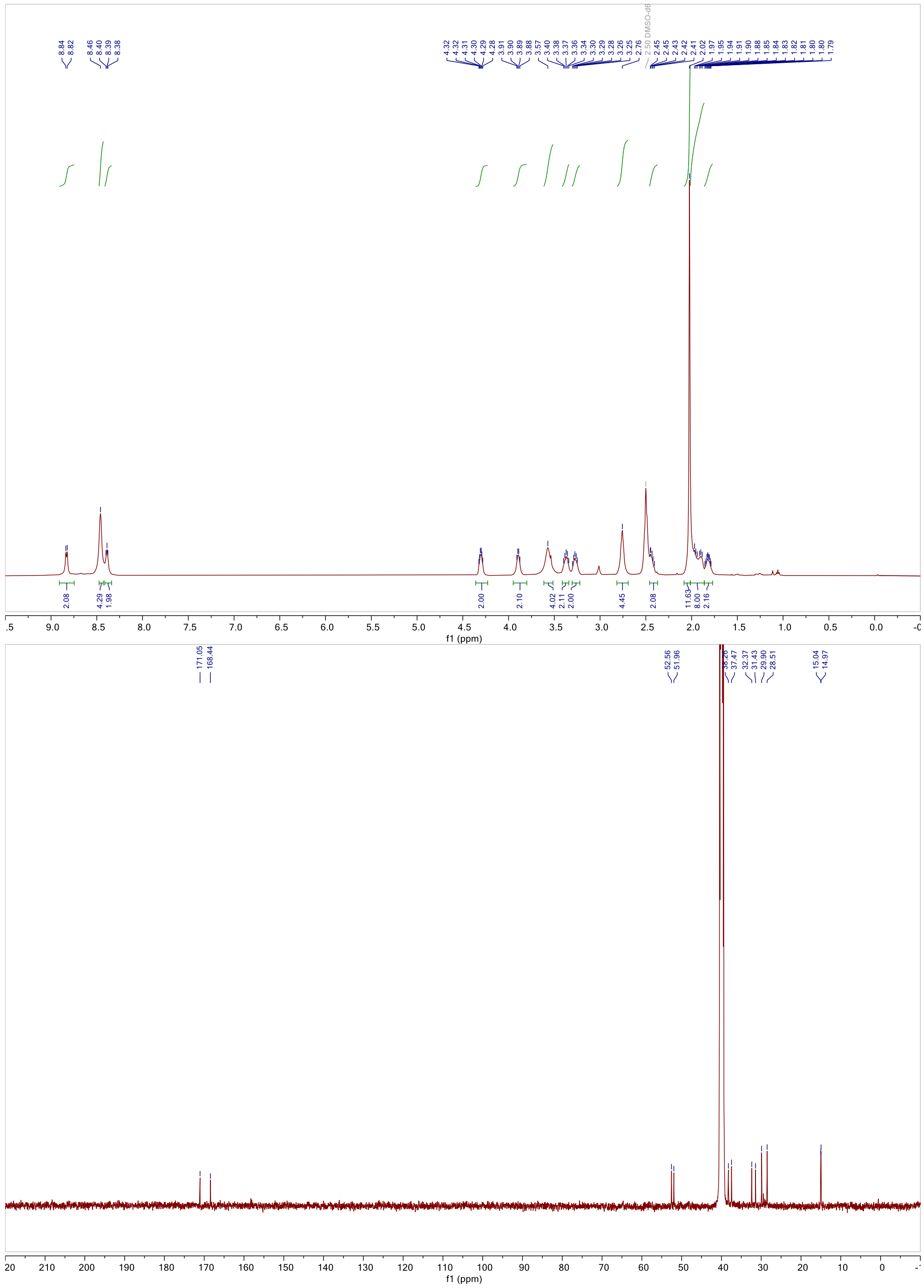


Fig. S14f $^{1}$H NMR (top) and $^{13}$C NMR (bottom) spectra of MMssMM in DMSO-$d_6$.

g. FFssFF (624 mg, 60.1 %); white power; **$^{1}$H NMR** (500 MHz, DMSO-$d_6$) $\delta$ 8.82 (d, $J$ = 8.1 Hz, 2H), 8.34 (t, $J$ = 5.7 Hz, 2H), 8.07 (s, 4H), 7.35–7.18 (m, 20H), 4.53 (td, $J$ = 8.1, 6.0 Hz, 2H), 4.04 (s, 2H), 3.29 (dq, $J$ = 13.2, 6.5 Hz, 4H), 3.11 (dd, $J$ = 14.2, 5.1 Hz, 2H), 2.96 (ddd, $J$ = 26.2, 13.9, 7.0 Hz, 4H), 2.86 (dd, $J$ = 13.7, 8.2 Hz, 2H), 2.68 (hept, $J$ = 6.7 Hz, 4H); **$^{13}$C NMR** (125 MHz, DMSO-$d_6$) $\delta$ 170.77, 168.22, 137.68, 135.15, 130.03, 129.63, 128.92, 128.65, 127.57, 126.94, 54.79, 53.54, 38.45, 38.40, 37.39, 37.34.

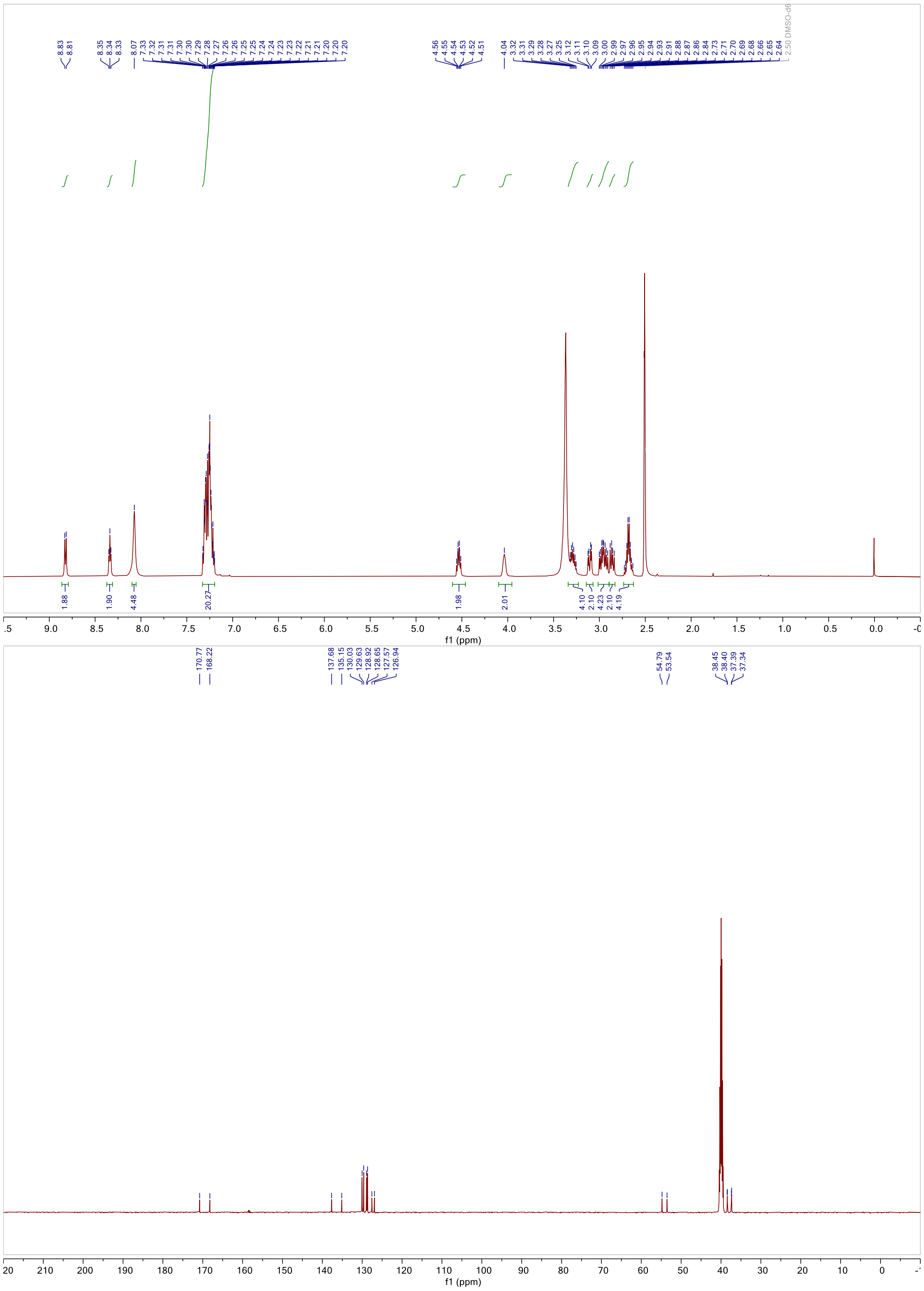


Fig. S14g $^{1}H$ NMR (top) and $^{13}C$ NMR (bottom) spectra of FFssFF in DMSO-$d_6$.

h. WWssWW (782 mg, 62.3%); white powder; **$^{1}$H NMR** (500 MHz, DMSO-$d_6$) $\delta$ 11.00 (d, $J$ = 2.4 Hz, 2H), 10.85 (d, $J$ = 2.4 Hz, 2H), 8.90 (d, $J$ = 7.9 Hz, 2H), 8.27 (t, $J$ = 5.7 Hz, 2H), 7.95 (d, $J$ = 5.3 Hz, 4H), 7.70 (d, $J$ = 7.9 Hz, 2H), 7.62 (d, $J$ = 7.9 Hz, 2H), 7.34 (dd, $J$ = 16.5, 8.1 Hz, 4H), 7.18 (dd, $J$ = 16.7, 2.4 Hz, 4H), 7.12–7.03 (m, 4H), 6.99 (dt, $J$ = 11.8, 7.4 Hz, 4H), 4.58 (q, $J$ = 7.4 Hz, 2H), 4.05–4.00 (m, 2H), 3.40–3.34 (m, 2H), 3.26 (dt, $J$ = 15.7, 4.2 Hz, 4H), 3.14 (dd, $J$ = 14.5, 6.2 Hz, 2H), 3.03 (dt, $J$ = 14.5, 9.0 Hz, 4H), 2.63 (hept, $J$ = 6.6 Hz, 4H); **$^{13}$C NMR** (125 MHz, DMSO-$d_6$) $\delta$ 171.38, 168.77, 136.78, 136.53, 127.71, 127.50, 125.55, 124.23, 121.61,

121.39, 119.03, 118.88, 118.70, 111.91, 111.78, 109.99, 107.15, 54.37, 52.82, 38.51, 37.20, 28.70, 27.95.

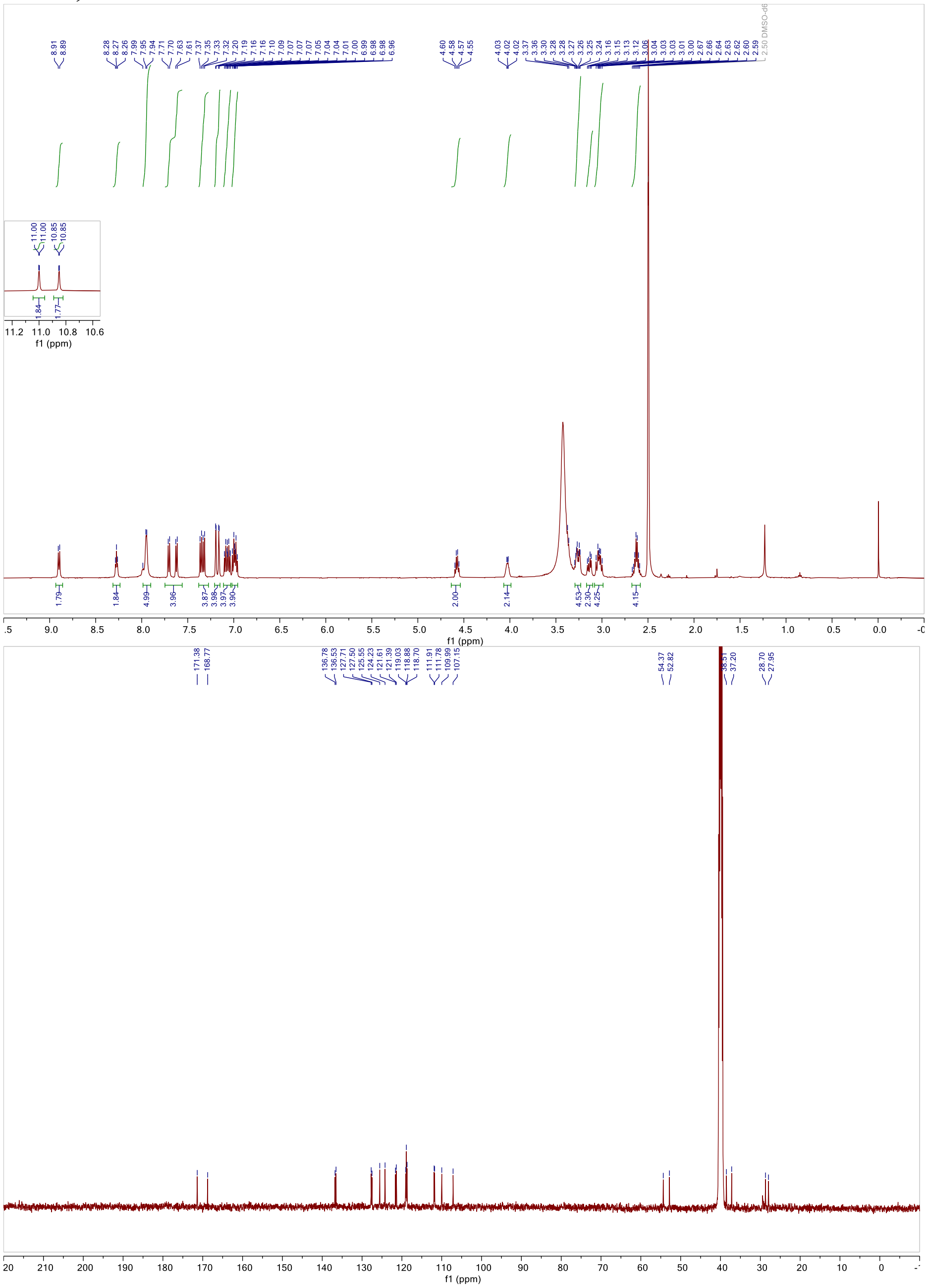


Fig. S14h $^{1}H$ NMR (top) and $^{13}C$ NMR (bottom) spectra of WWssWW in DMSO-$d_6$.

*ESI mass spectrometry of tetrapeptides*

a. AAssAA

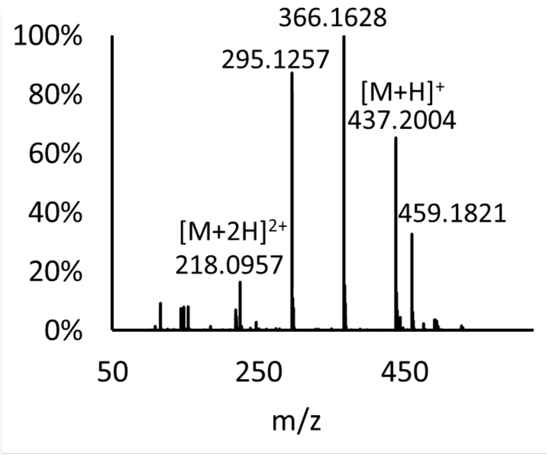


Fig. S15a Mass spectrum of AAssAA. [M+H]$^+$ calculated for $C_{16}H_{33}N_6O_4S_2$ 437.2005, found 437.2004.

b. PPssPP

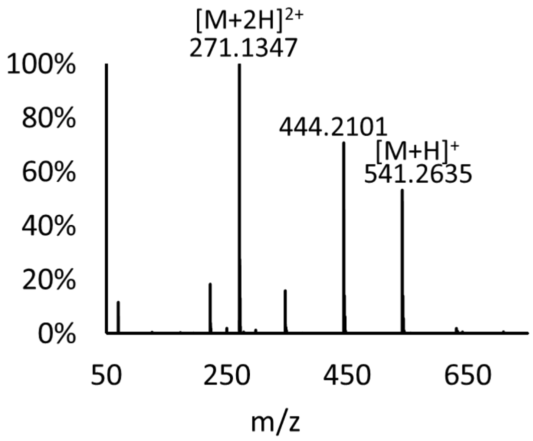


Fig. S15b Mass spectrum of PPssPP. [M+H]$^+$ calculated for $C_{24}H_{41}N_6O_4S_2$ 541.2631, found 541.2635.

c. VVssVV

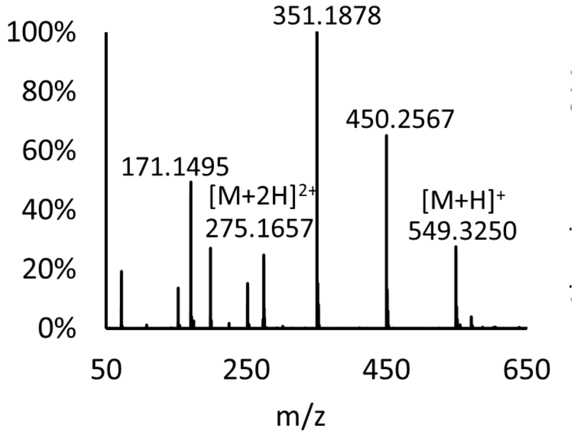

Fig. S15c Mass spectrum of VVssVV. $[M+H]^+$ calculated for $C_{24}H_{49}N_6O_4S_2$ 549.3257, found 549.3250.

d. LLssLL

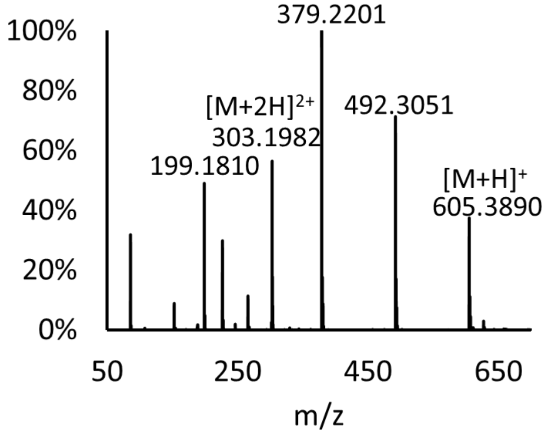


Fig. S15d Mass spectrum of LLssLL. $[M+H]^+$ calculated for $C_{28}H_{57}N_6O_4S_2$ 605.3883, found 605.3890.

e. IIssII

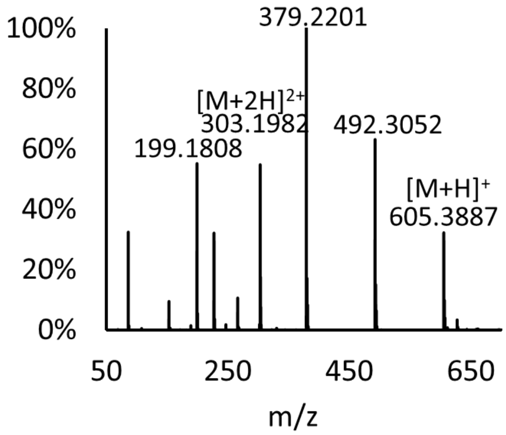


Fig. S15e Mass spectrum of IIssII. $[M+H]^+$ calculated for $C_{28}H_{57}N_6O_4S_2$ 605.3883, found 605.3887.

f. MMssMM

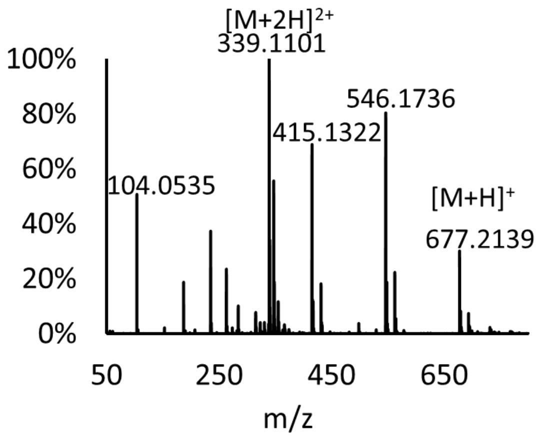


Fig. S15f Mass spectrum of MMssMM. $[M+H]^+$ calculated for $C_{24}H_{49}N_6O_4S_6$ 677.2140, found 677.2139.

g. FFssFF

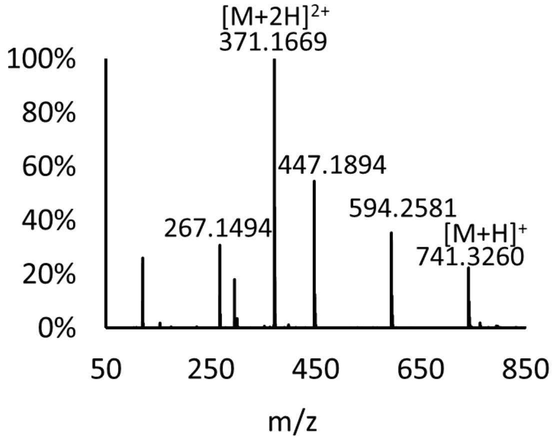


Fig. S15g Mass spectrum of FFssFF. $[M+H]^+$ calculated for $C_{40}H_{49}N_6O_4S_2$ 741.3257, found 741.3260.

h. WWssWW

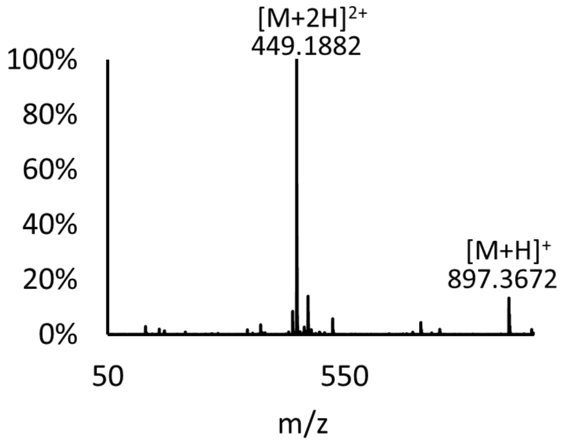

Fig. S15h Mass spectrum of WWssWW. $[M+H]^+$ calculated for $C_{48}H_{53}N_{10}O_4S_2$ 897.3693, found 897.3672.

*2. Dye synthesis, purification, and characterization*

**Materials.** 1-iodoctadecane, 4-diethylaminobenzaldehyde, and trimethylsilyl chloride were from Sigma Aldrich. 2-methybenzoxazole was from TCI Chemicals.

**Synthesis.** 2-methybenzoxazole (5.2 mmol, 1 equiv) and 1-iodoctadecane (6.24 mmol, 1.2 equiv) were dissolved in 15 mL acetonitrile and introduced into a round-bottom flask. The reaction mixture was stirred under nitrogen atmosphere at 85 °C until completion (typically 24 hours, monitored by TLC). The reaction mixture was concentrated in a rotary evaporator and washed with 70 mL of diethyl ether, and the purple precipitate was collected after centrifugation. The crude material (4.33 mmol, 1 equiv), 4-diethylaminobenzaldehyde (4.33 mmol, 1 equiv), and trimethylsilyl chloride (109.9 uL, 0.2 equiv) were dissolved in 8 mL DMF and refluxed at 135 °C overnight until completion (monitored by TLC). The reaction mixture was cooled to room temperature and extracted with dichloromethane (DCM)/$H_2O$ three times to extract the organic layer.

**Purification.** Purification was performed using column chromatography (DCM/MeOH, 15:1 ratio).

**Characterization.** $^1H$ and $^{13}C$ NMR spectra were recorded on a Bruker AV-500 spectrometer.

*NMR spectra of tetrapeptides*

Dark green solid (1236.2 mg, 49.95%), **$^1$H NMR** (500 MHz, DMSO-$d_6$) $\delta$ 8.35 (d, $J$ = 15.4 Hz, 1H), 8.08 (d, $J$ = 8.8 Hz, 2H), 7.80 (d, $J$ = 7.4 Hz, 1H), 7.73 (d, $J$ = 8.0 Hz, 1H), 7.56 (t, $J$ = 7.0 Hz, 1H), 7.49 (t, $J$ = 7.2 Hz, 1H), 7.24 (d, $J$ = 15.6 Hz, 1H), 6.90 (d, $J$ = 9.4 Hz, 2H), 4.56 – 4.46 (m, 2H), 3.59 (m, $J$ = 7.1 Hz, 4H), 1.77 (m, 8H), 1.43 – 1.20 (m, 36H), 0.90 – 0.85 (t, 3H); **$^{13}$C NMR** (125 MHz, DMSO-$d_6$) $\delta$ 179.57, 154.76, 153.14, 143.05, 141.54, 129.24, 127.83, 123.24, 122.46, 113.96, 112.47, 104.61, 51.25, 45.27, 44.92, 31.72, 29.41, 29.27, 29.24, 29.11, 28.99, 28.07, 27.03, 26.19, 22.52, 14.38, 12.99.

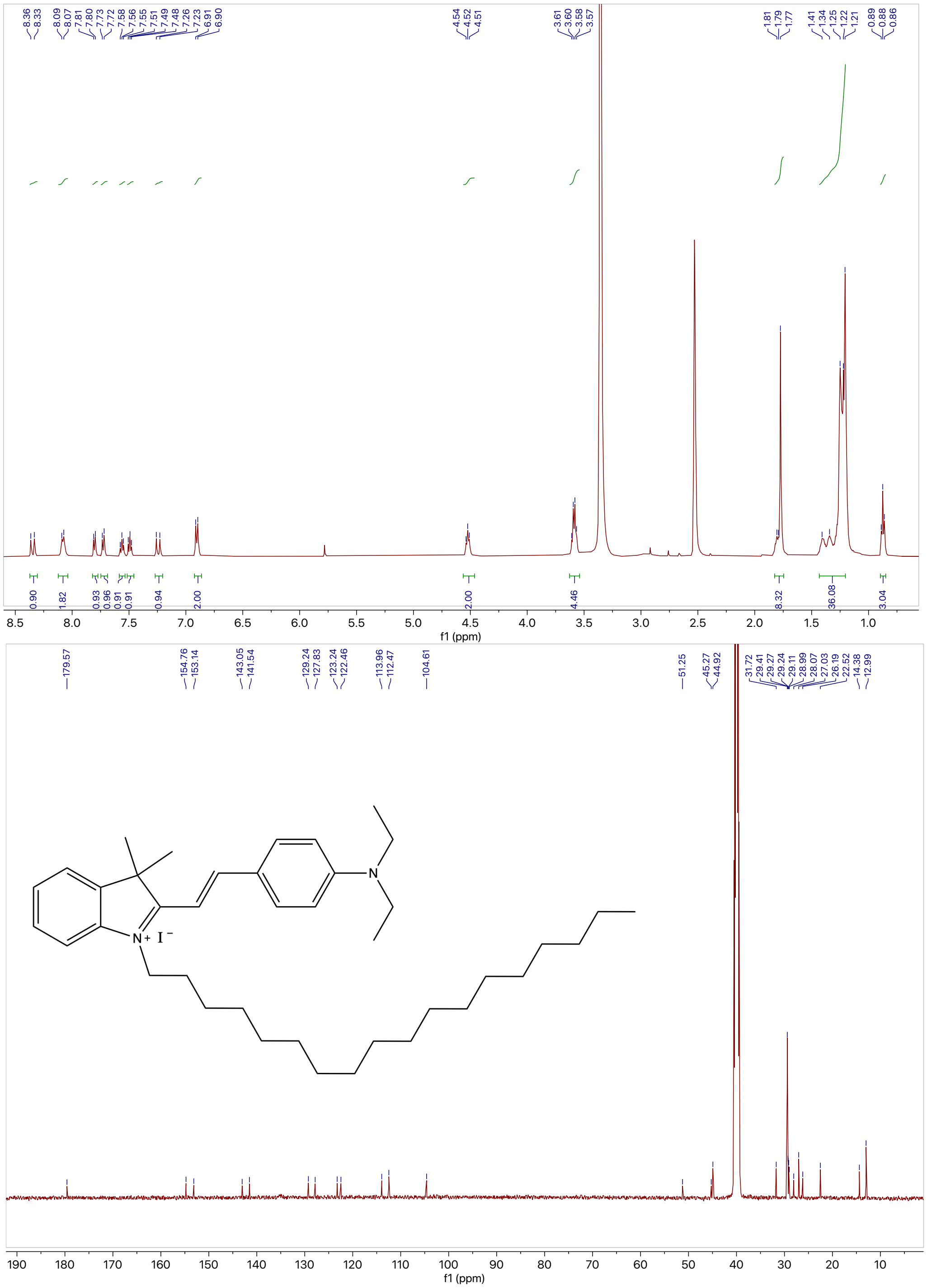


Fig. S16 $^{1}H$ NMR (top) and $^{13}C$ NMR (bottom) spectra of the dye molecule in DMSO-$d_6$. Inset: chemical structure of the dye molecule.
